# DACO: Distortion/artefact correction for diffusion MRI data in an integrated framework

**DOI:** 10.1101/2021.07.06.450481

**Authors:** Yung-Chin Hsu, Wen-Yih Isaac Tseng

## Abstract

In this paper we propose a registration-based algorithm to <u>co</u>rrect various <u>d</u>istortions or <u>a</u>rtefacts (DACO) commonly observed in diffusion weighted (DW) magnetic resonance images (MRI). The registration in DACO is proceeded on the basis of a pseudo *b*_0_ image, which is synthesized from the anatomical images such as T1-weighted image or T2-weighted image, and a pseudo diffusion MRI (dMRI) data, which is derived from the Gaussian model of diffusion tensor imaging (DTI) or the Hermite model of MAP-MRI. DACO corrects (1) the susceptibility-induced distortions, (2) the intensity inhomogeneity, and (3) the misalignment between the dMRI data and anatomical images by registering the real *b*_0_ image to the pseudo *b*_0_ image, and corrects (4) the eddy current (EC)-induced distortions and (5) the head motions by registering each of the DW images in the real dMRI data to the corresponding image in the pseudo dMRI data. As the above artefacts interact with each other, DACO models each type of artefact in an integrated framework and estimates these models in an interleaved and iterative manner. The mathematical formulation of the models and the comprehensive estimation procedures are detailed in this paper. The evaluation using the human connectome project data shows that DACO could estimate the model parameters accurately. Furthermore, the evaluation conducted on the real human data acquired from clinical MRI scanners reveals that the method could reduce the artefacts effectively. The DACO method leverages the anatomical image, which is routinely acquired in clinical practice, to correct the artefacts, minimizing the additional acquisitions needed to conduct the algorithm. Therefore, our method is beneficial to most dMRI data, particularly to those without acquiring the field map or blip-up and blip-down images.

## 1. Introduction

Diffusion magnetic resonance imaging (dMRI) is a unique tool to explore the microstructural architectures of brain tissues and has been used in various clinical applications. This technique uses a pair of diffusion-sensitive gradients to encode the Brownian motion of water molecules into the intensity of the diffusion-weighted (DW) images. To recover the microstructural information encoded in the DW signals, a dMRI data usually comprises a set of DW images acquired with gradients with various magnitudes (the b-values) and along various orientations (the b-vectors). Many algorithms, such as diffusion tensor imaging (DTI) (Basser et al., 1994) and MAP-MRI (Özarslan et al., 2013), have been proposed to reconstruct average diffusional propagators of the ensembles of water molecules from the dMRI data.

Although dMRI is a powerful tool, dMRI data is corrupted by various artefacts. To make the acquisition time practical, the echo-planar imaging (EPI) technique is often used to accelerate the speed of acquisition. However, EPI is sensitive to off-resonance deviations, rendering DW images suffer from two types of spatial distortions. The first one is the susceptibility-induced distortions (Andersson et al., 2003; Hutton et al., 2002), which originate from the inhomogeneous B0 fields (the so-called field map) due to the presence of abrupt susceptibility difference between the brain and neighboring tissues. The other is the eddy current (EC)-induced distortions (Jezzard et al., 1998), which are caused by the residual effects of diffusion-sensitive gradients. Since in the blipped EPI sequence the acquisition bandwidth of phase-encoding (PE) is substantially lower than that of frequency-encoding (FE), the distortions are much more pronounced along the PE direction than the FE direction (Hutton et al., 2002). Therefore, the distortions are usually assumed to occur along the PE direction (Andersson et al., 2003; Irfanoglu et al., 2015), simplifying the problem to one dimension (1D). Besides the distortion artefacts, DW images usually do not align to each other because of inevitable head motions during long acquisition of the dMRI data. Another artefact usually observed in the dMRI data is the intensity inhomogeneity (sometimes called bias field) (McVeigh et al., 1986). This well-known artefact originates from the hardware characteristics, such as the imperfect B1 field transmission and the non-uniform sensitivity of receiving coils (McVeigh et al., 1986). These artefacts could prevent the dMRI processing algorithms from accurately extracting the structural information from the dMRI data, such as fractional anisotropy (FA) (Basser & Pierpaoli, 1996) or trajectories of tract bundles (Basser et al., 2000; Yeh et al., 2013). Another issue usually encountered in practice is the rigid alignment between the anatomical images, such as T1-weighted (T1w) image or T2-weighted (T2w) image, and the dMRI data. Obviously, a good alignment could facilitate the integration of the information provided by the anatomical images and the dMRI data, but a good alignment could only be achieved when the artefacts are properly reduced.

Various methods have been proposed to correct the aforementioned artefacts of EPI. For the susceptibility-induced distortions, most methods correct the artefact retrospectively by measuring or estimating the field map. The algorithms could be roughly divided into three categories. The first is field map-based, this method explicitly measures the field map by acquiring two gradient echo images with exact scanning parameters except for the echo time (TE), then the field map is the phase differences between the two images divided by the differences of the TE values. The second is reverse gradients-based (Andersson et al., 2003; Hedouin et al., 2017; Irfanoglu et al., 2015), where two *b*_0_ images (or two complete dMRI datasets) with reverse PE gradients are acquired. The *b*_0_ images here refer to the DW images whose b-value=0 s/mm^2^, i.e., without applying the diffusion-sensitive gradients. The basic idea behind this category is that the two *b*_0_ images (or the two dMRI datasets) would have reverse distortions, hence the field map could be estimated by registering them to their “midpoint” image, which is theoretically the undistorted image. The last is anatomical image-based (Bhushan et al., 2015; Huang et al., 2008), in this category, the real *b*_0_ image is registered to the pseudo *b*_0_ image, which is constructed from the anatomical images, such as T1w image or T2w image. Since the anatomical images are free of distortions, the estimated displacement map of the registration could be considered as a surrogate of the field map. Technically, both the reverse gradients-based and the anatomical image-based methods greatly involve registration algorithms, therefore, some efforts have been made to combine them by regarding the pseudo *b*_0_ images as the midpoint images in the reverse gradients-based methods; this approach has been shown to improve the performance of distortion correction (Irfanoglu et al., 2015).

The EC-induced distortions could be prospectively reduced through dedicated pulse sequences (Reese et al., 2003), however, the longer TE and longer repetition time (TR) would lead to lower signal-to-noise ratio (SNR) and longer acquisition time, both are detrimental to dMRI. Alternatively, the distortions could be reduced retrospectively by registering each of the DW images to the *b*_0_ image (Jezzard et al., 1998), which is usually assumed not corrupted by the EC-induced distortions. Another feasible way is to synthesize dMRI dada, in which the DW images are believed to have less EC-induced distortions, and the correction could be conducted by registering the raw DW images to the synthesized DW images (Andersson & Sotiropoulos, 2016). Similar retrospective strategies for the EC-induced distortions are also applicable to head motions, therefore, many algorithms conduct the correction for both artefacts altogether (Andersson & Sotiropoulos, 2016).

The intensity inhomogeneity is usually less concerned in the artefact correction algorithms, therefore, if this artefact is to be corrected in a clinical study, it is conducted after other artefacts have been corrected. The correction is through explicitly measuring the bias field (Wicks et al., 1993) or estimating the bias field using post-processing algorithms (Ashburner & Friston, 2005; Sled et al., 1998). Similarly, the alignment between the anatomical image and the dMRI data is often conducted separately, either before or after the artefact correction is performed (Glasser et al., 2013). Note that this alignment could be integrated with the correction of the susceptibility-induced distortions in the anatomical image-based methods (Bhushan et al., 2015).

In this study we propose the DACO method, which is a registration-based algorithm designed to <u>co</u>rrect the <u>d</u>istortions or <u>a</u>rtefacts of the dMRI data. To make the registration work, the method synthesizes a pseudo *b*_0_ image, which is constructed from the anatomical MRI image (T1w or T2w) using histogram equalization (Gonzalez & R.E., 2001), and a pseudo dMRI data, which is derived from the Gaussian model (i.e., DTI) or the Hermite model (i.e., MAP-MRI). DACO addresses five dMRI artefacts in an integrated framework. Explicitly, it uses the pseudo *b*_0_ image to drive (1) the correction for the susceptibility-induced distortions, (2) the correction for the intensity inhomogeneity, and (3) the spatial rigid misalignment between the anatomical image and the dMRI data, and uses the pseudo dMRI data to drive (4) the correction for the EC-induced distortions and (5) the correction for the head motions. To the best of our knowledge, DACO is the first to address these artefacts altogether in an integrated framework. The aim of this study is to describe the DACO method and to demonstrate the effectiveness and efficiency of the method in correcting dMRI artefacts.

## 2. Theory

Figure 1 illustrates the schematic diagram of the DACO algorithm. Two MRI data are needed in the registration process, the anatomical image (usually T1w or T2w image) for constructing the pseudo *b*_0_ image, and the dMRI data on which the artefact correction is applied. The pseudo *b*_0_ image, synthesized either from the T1w or T2w image, is denoted as 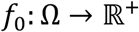, where 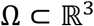 denotes the image domain. The voxel-coordinate ***x***_*f*_0__ (a 3×1 vector) of *f*_0_ is mapped to the millimeter (mm)-coordinate ***r***_*f*_0__ ∈ Ω through ***r***_*f*_0__ = ***M***_*f*_0__ ***x***_*f*_0__ + ***T***_*f*_0__, where ***M***_*f*_0__ is a 3×3 matrix comprising the information of the orientation and voxel size of *f*_0_, and ***T***_*f*_0__ a 3×1 vector denoting translation. Since the dMRI data comprises *n_d_* real DW images 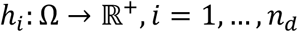, and each *h_i_* comes with a b-value 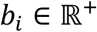 and a b-vector 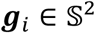, these three variables are expressed as (*h_i_, b_i_*, ***g***_*i*_). Here, *h*_0_ denotes the first raw *b*_0_ image of the dMRI data. In the study, we assume that *h_i_* are acquired with the same EPI parameters, including the same TE and TR, the same direction of PE 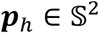 and echo spacing, the same numbers of voxels along the three spatial dimensions, and the same mapping matrix ***M***_*h*_ and translational vector ***T***_*h*_ such that the voxel-coordinate ***x***_*h_i_*_ (a 3×1 vector) of *h_i_* is mapped to the mm-coordinate ***r***_*h_i_*_ ∈ Ω by ***r***_*h*_ = ***M***_*h*_***x***_*h_i_*_ + ***T***_*h*_.

**Fig. 1.**
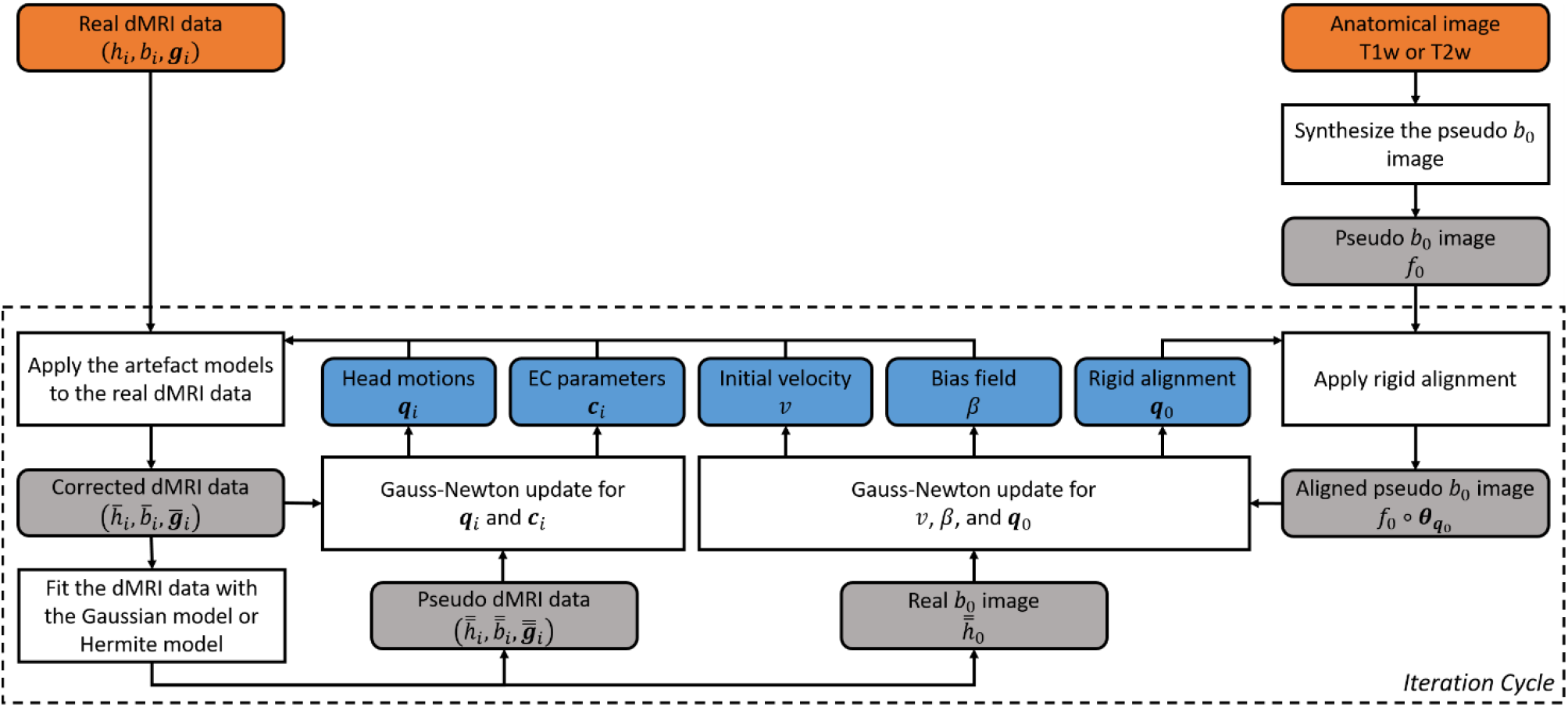
The schematic diagram of the DACO algorithm.

To estimate the parameters of the artefact models in the same integrated framework, the estimation is conducted in an iterative manner (see Fig. 1). In every iteration, the estimated parameters of the artefact models are applied on the raw dMRI data to create the corrected data, denoted as 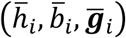. The pseudo dMRI data are also constructed in every iteration by fitting 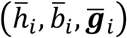 with the Gaussian or Hermite model, they are denoted as 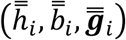; particularly, the image whose b-value is zero is constructed and denoted as 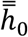. In the DACO method, the natural choice of the reference space during the entire estimation process is the space of 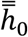, therefore, all of the derived images or maps are situated in this space. By design, we specifically let the space of 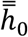 align with the space of the first DW image (i.e., *h*_1_).

### 2.1 Illustration of Artefact Models

In this section, we illustrate the synthesis procedures of the pseudo *b*_0_ image and the pseudo dMRI data as well as the mathematical formulation for each artefact model. The optimization algorithm is described in the next section by putting these artefact models altogether.

#### 2.1.1 Synthesis of Pseudo *b*_0_ Image

Knowing that most dMRI sequences use the pulsed gradient spin echo (PGSE) technique, rendering the contrast of the real *b*_0_ images inherently T2-weighted, two approaches have been proposed to synthesize the pseudo *b*_0_ image. The first approach directly uses the conventional T2w image for the construction (Huang et al., 2008). The second approach synthesizes the pseudo *b*_0_ image by inverting the contrast of T1w image (Bhushan et al., 2015) based on the fact that the intensity order of the three major brain tissues, i.e. white matter (WM), gray matter (GM), and cerebrospinal fluid (CSF), of the T1w image and that of the T2w / *b*_0_ image are roughly reversed. The construction procedures are described as follow.

1. Create a brain mask which comprises the WM, GM, and CSF regions from the T1w or T2w image.
2. Correct the intensity inhomogeneity of the T1w or T2w image using an externally measured bias field (Wicks et al., 1993) or through post-processing algorithms (Ashburner & Friston, 2005; Sled et al., 1998).
3. Apply the brain mask of (1) on the result of (2).
4. For T1w image, apply contrast inversion to the result of (3), for T2w image, do nothing.
5. Conduct histogram equalization with respect to the real *b*_0_ image on the result of (4). Since the problem of the intensity inhomogeneity on the T1w or T2w image has been corrected in the procedures, the synthesized pseudo *b*_0_ image is distortion-free and of little inhomogeneity problem.

#### 2.1.2 Rigid Alignment between Real and Pseudo *b*_0_ Images

Several factors contribute to the image discrepancy between the pseudo and real *b*_0_ images. The first factor is the rigid misalignment (i.e., the real *b*_0_ image does not align with the pseudo *b*_0_ image), which may happen during a long acquisition period. The spatial misalignment could be represented by ***θ***_*q*_0__:Ω → Ω with 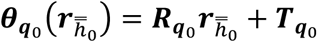 such that 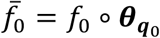 is in alignment with 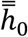, where ∘ denotes function composition. Here ***q***_0_ is a 6×1 vector (with elements *q_0j_, j* = 1…6) which comprises three elements representing translation (***T***_*q*_0__) and three Euler angles parameterizing rotation (***R***_*q*_0__). The spatial misalignment can be represented in the inverted way such that 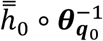 aligns with *f*_0_, where 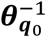 is the inverse of ***θ***_*q*_0__.

#### 2.1.3 Intensity Bias

Another factor of the image discrepancy is the intensity bias, intrinsic and extrinsic, of the real *b*_0_ image with respect to the pseudo *b*_0_ image. The extrinsic intensity bias originates from the intensity inhomogeneity of the real *b*_0_image, and can be corrected using externally measured bias fields or through post-processing algorithms, as those applied on the raw T1w or T2w image. However, the presence of the susceptibility-induced distortions in the real *b*_0_ image might hinder the performance of correction. Another source of the intensity bias arises from the fact that the real and pseudo *b*_0_ images are obtained from different modalities. Such bias is inherent and cannot be fully removed by histogram equalization. In the study, we consider the intrinsic and extrinsic intensity biases altogether and extend the meaning of “bias field” to represent the collective effects. We assume that the bias field is multiplicative and model it in terms of *e^β^*, where 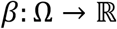. In this way, the contrast of 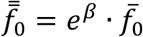 would be similar to the real *b*_0_ image. Practically, the *β* field is assumed to be sufficiently smooth, in order to avoid the situation that the intensity bias explains most of the differences between the real and pseudo *b*_0_ images, which is certainly not the desired results. In the study, the smoothness of the *β* field is achieved via the differential operator *L_β_*, with the norm of *β* defined as ||*β*||^2^ = 〈*L_β_β*|*β*〉_2_.

#### 2.1.4 Susceptibility-induced Distortions

The last confounding factor of the image discrepancy is the susceptibility-induced distortions. This factor induces the spatial discrepancy between the distortion-free (the pseudo *b*_0_ image) and the distorted image (the real *b*_0_ image). We assume that the distortion map ***φ***:Ω → Ω belongs to Diff(Ω), the group of diffeomorphisms (i.e., one-to-one, reversible, and smooth transformations that preserve topology), hence by definition ***φ***^−1^, the inverse of ***φ***, exists and also belongs to Diff(Ω). Furthermore, we assume that an image *f* is distorted by ***φ*** according to the action ***φ*** * *f* = |D***φ***^−1^| · *f* ∘ ***φ***^−1^, where D is the Jacobian operator and |·| is the determinant. This assumption has been widely employed in various algorithms aiming at correcting the susceptibility-induced distortions (Andersson et al., 2003; Bhushan et al., 2015; Bhushan et al., 2012; Hedouin et al., 2017; Irfanoglu et al., 2015).

The diffeomorphisms can be constructed under the framework of Large Deformation Diffeomorphic Metric Mapping (LDDMM) (Beg et al., 2005; Miller et al., 2006) by shooting the initial velocity field 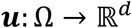, *d* ∈ {1,2,3} to generate a series of velocity fields 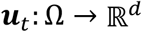, *t* ∈ [0,1] with ***u***_0_ = ***u*** and construct the flow of diffeomorphisms ***φ***_*t*_:Ω → Ω with ***φ***_0_ = ***Id*** (i.e., the identity). More specifically, we solve the system of equations

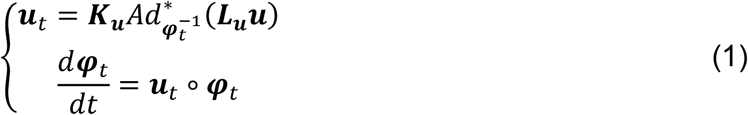

where the adjoint operator is defined as 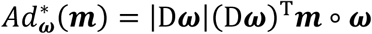, and ^T^denotes matrix transpose. To ensure that the constructed deformations are diffeomorphisms, the velocities ***u***_*t*_ have to be sufficiently smooth; the smoothness is controlled by the differential operator ***L***_*u*_ and its Green’s function ***K***_*u*_. Under the LDDMM framework, the original problem of searching for the distortion map ***φ*** turns out to the problem of estimating the initial velocity ***u*** such that ***φ***_1_ = ***φ***. The norm of ***u***_*t*_ is defined through ||***u***_*t*_||^2^ = 〈***L***_*u*_***u***_*t*_|***u***_*t*_〉_2_.

In the blipped EPI sequence, it is appropriate to consider the distortions along the PE direction only (Hutton et al., 2002), reducing the susceptibility-induced distortions to the 1D-LDDMM (i.e., *d* = 1) problem, one for each PE column (with the image domain Ω_*p*_ ⊂ Ω). In this manner, the velocities are functions of 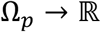 and diffeomorphisms are functions of Ω_*p*_ → Ω_*p*_. From the theoretical perspective, these 1D-LDDMM problems can be solved independently, and so the initial velocities and the diffeomorphisms of the PE columns are independent. Although the 1D formulation ensures the smoothness along each PE column, it does not ensure the smoothness between adjacent columns. This makes the deformation zigzag across the columns, as shown previously (Andersson et al., 2003). To mitigate the problem, we slightly modify the 1D-LDDMM formulation by considering these 1D problems together rather than independently. More precisely, we gather the initial velocities column by column to form a 3D collective initial velocity field 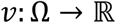, and use a 3D differential operator *L_v_* (and its Green’s function *K_v_*) to ensure the smoothness along and across the PE columns. The norm of *v* is defined through ||*v*||^2^ = 〈*L_v_v*|*v*〉_2_. Except this modification, the 1D-LDDMM formulation remains unchanged. For each PE column, the displacements induced by the distortions could be obtained by subtracting the identity from the deformations at *t* = 1. Like the initial velocities, we could collect the displacements column by column to form a displacement map. Because *v* is sufficiently smooth, the displacement map is also smooth along and across the PE columns. Theoretically, the displacements are proportional to the off-resonance frequencies (i.e., the field map) induced by the susceptibility under the linear approximation. In the study, we use 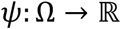 to denote the field map (in unit of Hertz). The relation between the distortion map and the field map is

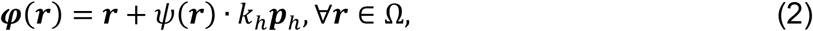

where *k_h_* = *τV, τ* is the effective echo-spacing time and *V* is the field of view (FOV) of the EPI image in the PE direction. In addition to the real *b*_0_ image, other DW images are also distorted in the same fashion, except that the field map is the rotated version accompanied with the head motion. This issue will be discussed in detail.

#### 2.1.5 Synthesis of Pseudo dMRI Data

In each iteration, we synthesize a pseudo dMRI data to update the parameters for the EC-induced distortions and head motions. In this manner, a series of updated pseudo dMRI data having less and less artefact are constructed and the parameters could gradually converge to an optimal solution. For dMRI data where the maximum b-value (b-max) is smaller than 1500 s/mm^2^, the pseudo dMRI data is synthesized through the Gaussian model of DTI (Basser et al., 1994). In contrast, if b-max exceeds 1500 s/mm^2^, the Hermite model of MAP-MRI (Özarslan et al., 2013) is employed instead. While DTI fits the diffusion signals with an anisotropic Gaussian function, MAP-MRI models the diffusion signals with a linear combination of Hermite functions. MAP-MRI is a generalization of DTI, since the Hermite function of zeroth order is a Gaussian function (Özarslan et al., 2013). In the study, the maximum order of the Hermite functions is four (Ning et al., 2015). Put it more precisely, the construction procedures are as follow.

1. Apply the artefact models estimated in the previous iteration, including the susceptibility-induced distortions, intensity biases, EC-induced distortions, and head motions, on the raw dMRI data, to produce the corrected dMRI data 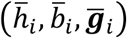
2. Fit the Gaussian model of DTI (b-max<1500 s/mm^2^) or the Hermite model of MAP-MRI (b-max≧1500 s/mm^2^) to 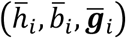 Results of this step are the diffusion tensors or the MAP-MRI coefficients.
3. Use the Gaussian model or the Hermite model to construct the pseudo dMRI data 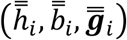 and the real *b*_0_ image 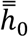 in accordance with the results of (2).

#### 2.1.6 Head Motions

The spatial discrepancy between *h_i_* and 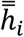 induced by head motions, like the rigid misalignment between the real and pseudo *b*_0_ images, is expressed by ***θ***_*q_i_*_:Ω → Ω with 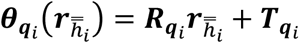 such that *h_i_* ∘ ***θ***_*q_i_*_ is in alignment with 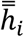. Here, ***q***_*t*_ are 6×1 vectors (with elements *q_ij_*, *j* = 1…6), each comprises three elements representing translation (***T***_*q*_) and three Euler angles parameterizing rotation (***R***_*q_i_*_). Recall that we specifically make the reference space to align with the space of *h*_1_, therefore, the estimated ***q***_*i*_ are rearranged in every iteration such that ***q***_1_ = **0**.

#### 2.1.7 Eddy Current-induced Distortions

Like the susceptibility-induced distortions, it is appropriate to consider the EC-induced distortions along the PE direction only. Let 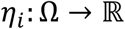 be the EC-induced magnetic off-resonance fields (in unit of Hertz) associated with *h_i_*. In the DACO method, *η_i_* are expressed as a linear combination of polynomial functions 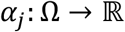 (Andersson & Sotiropoulos, 2016),

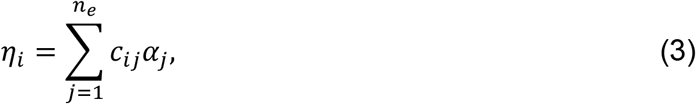

therefore, each *η_i_* is parameterized by a coefficient vector ***c**_i_* = [*c_i_1__,…, c_n_e__*]^*T*^. Let 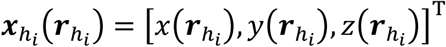 be the voxel-coordinate of point ***r***_*h_i_*_ and 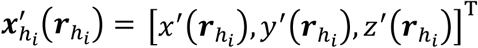 such that 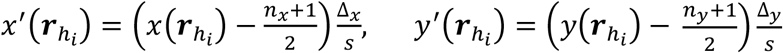 and 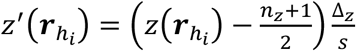, where *n*_{*x,y,z*}_ denotes the number of voxels and Δ_{*x,y,z*}_ denotes the voxel size (in unit of mm) along the {*x, y, z*} dimension, respectively. Here *s* is a scaling factor, and we set *s* = 100 mm in the study. The number of coefficients *n_e_* depends on the maximum order of the polynomials. For the linear model, *n_e_* = 4 and {*α_j_|j* = 1,2,3,4} = {*x*′, *y*′, *z*′, 1}. For the quadratic model, *n_e_* = 10 and 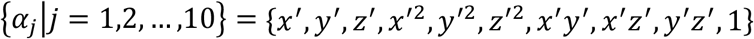. Similar to the susceptibility-induced distortions, the relation between the EC-induced distortions and the EC-induced off-resonance field is 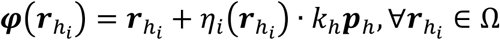.

#### 2.1.8 Combination of Artefact Models

If all the artefact models are applied on the pseudo *b*_0_ image, the results would be

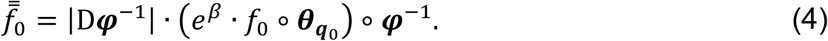

Here the pseudo *b*_0_ image is aligned to the space of the real *b*_0_ image by rigidlytransforming with ***θ***_*q*_0__, then multiplied with the bias field *e^β^* to make the appearance more like the real *b*_0_ image, and finally distorted by the distortion map ***φ***. For the DW images, the situation is more complicated because now we have to consider two more artefacts: the EC-induced distortions and head motions. In the DACO method, we assume that the field map is fixed to the head, therefore, when the head moves, the field map would be moved accordingly. More specifically, the field map *ψ* situates in the space of 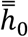, hence the field maps aligned with *h_i_* would be 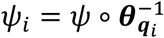. Together with the EC-induced off-resonance fields *η_i_*, which situate in the space of *h_i_*, the distortion map would be

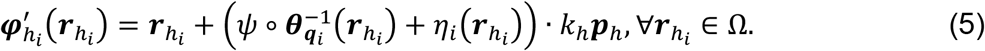

Expressing the distortion map in the space of 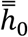 yields

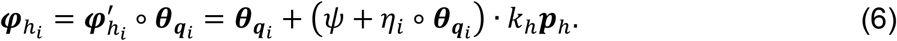

Clearly, ***φ***_*h_i_*_ are functions of ***q***_*i*_ and ***c***_*i*_. Applying the artefact models on 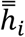 yields 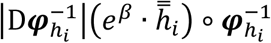. Conversely, the corrected dMRI data would be

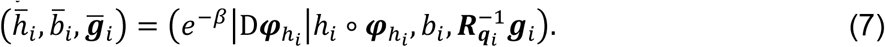

Because of the correction on head motions, the orientation of the diffusion-sensitive gradients should be rotated accordingly, but the b-values are assumed to be unchanged.

### 2.2 Estimation

In the DACO method, two energy functionals are used, one for the *b*_0_ images (*E*_1_) and another for the dMRI data (*E*_2_). We use the sum of squared differences (SSD) to quantify the data matching between real and pseudo *b*_0_ images, with the regularizations on the bias field and the initial velocity. Therefore, the data matching functional *E_d_* is

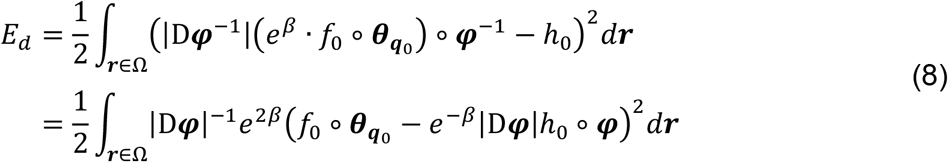

In Eq. (8), SSD is evaluated in the undistorted space, through change of variables. Sometimes, we use 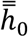 to approximate *e_−β_*|D***φ***|*h*_0_ ∘ ***φ*** because the two images have very similar appearance. In this way, *E_d_* can be evaluated as

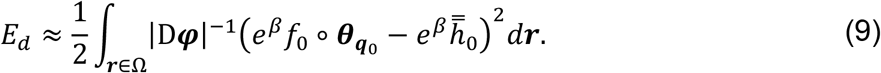

Finally, the total energy for *b*_0_ images is

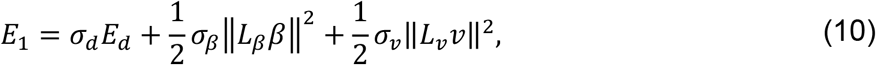

where *σ_d_, σ_β_*, and *σ_v_* are the weightings for the data matching and the regularization terms, respectively. For the dMRI data, we also employ SSD to evaluate the data matching. The energy functional *E*_2_ is

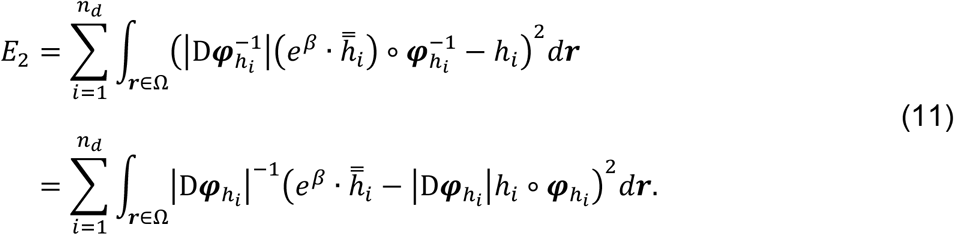

The goals of the two functionals are different. In every iteration, we update ***q***_0_, *β*, and *v* to reduce the *E*_1_ functional, then use the latest values of ***q***_0_, *β*, and *v* to calculate 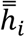 and update ***q***_*i*_ and ***c***_*i*_ to reduce the *E*_2_ functional.

For convenience, we define some assistant functions. The function ***b*** = vec(*A*) converts the 3D field 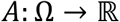 to the vector 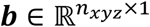 by concatenating the field column by column, where *n_xyz_* is the total number of voxels. The reverse operation is achieved by *A* = ivec(***b***). In addition, the function ***B*** = diag(***b***) converts the ***b*** vector to the diagonal matrix ***B***.

#### 2.2.1 Optimization of *q*_0_

We use the Gauss-Newton approach (Press & Numerical Recipes, 2007) to update the values of ***q***_0_. Let

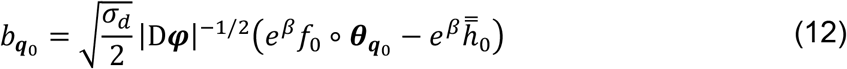

and

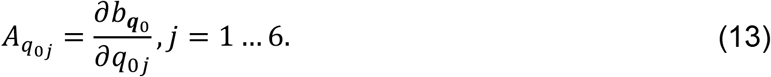

It follows that ***q***_0_ is updated by

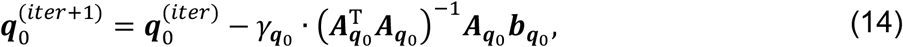

where 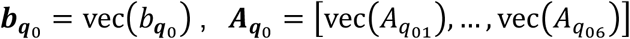, and 0 < *γ_q_0__* ≤ 1 is the scaling factor controlling the descent size of each iteration.

#### 2.2.2 Optimization of *β*

The first Gâteaux variation of *E_d_* with respect to *β* in the direction *ξ* is computed via

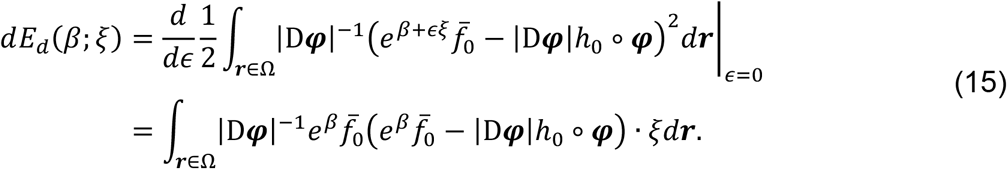

The second Gâteaux variation of *E_d_* with respect to *β* in the directions *ξ*_1_ and *ξ*_2_ is

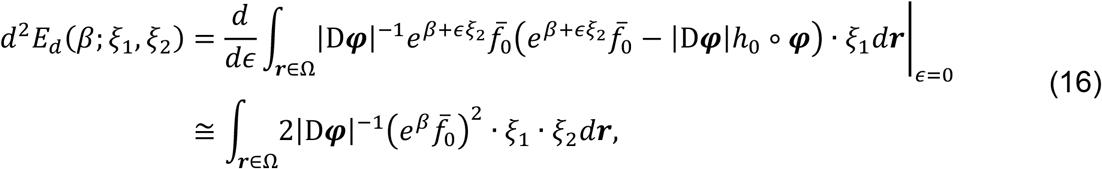

where the approximation is to make the second derivative positive definite. We employ the Gauss–Newton approach similar to (Ashburner & Friston, 2011) to update the values of *β*. Let

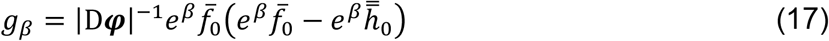

and

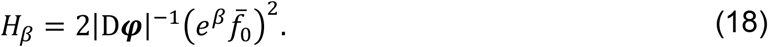

Also let ***b***_*β*_ = vec(*β*), ***g***_*β*_ = vec(*g_β_*), and ***H***_*β*_ = diag(vec(*H_β_*)). The descent deviation is computed via

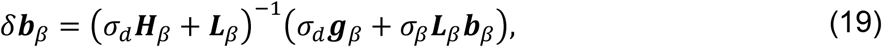

or in the 3D field form *δβ* = ivec(*δ**b**_β_*), then the *β* field is updated by

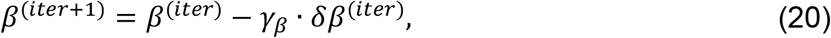

where 0 < *γ_β_* ≤ 1. Eq. (19) involves the inversion of a huge matrix, which is solved by the full multigrid algorithm as proposed in (Ashburner, 2007). In this study, *L_β_* is implemented through

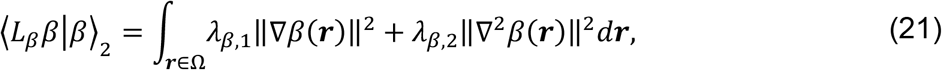

where the first term in the integral is the membrane energy and the second term is the bending energy, *λ*_*β*,1_ and *λ*_*β*,2_ control their strength of regularization, respectively. Practically, this differential operator is expressed in the matrix form 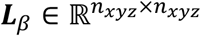.

#### 2.2.3 Optimization of *v*

In the Appendix A, we derive the first and second derivatives of *E_d_* with respect to the initial velocity ***u*** in Eq. (A13) and Eq. (A14), respectively. Recall that we modify the 1D LDDMM formulation by gathering the initial velocities of the PE columns to form the 3D collective initial velocity field *v*. In the same vein, the first and second derivatives of *E_d_* with respect to *v* are

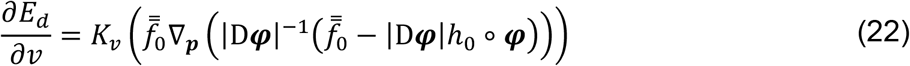

and

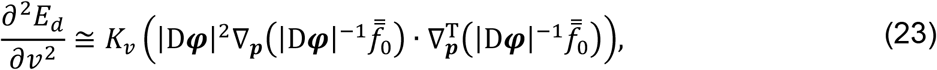

respectively. Though *v* is a scalar field, it is actually along the PE direction ***p***_*h*_, therefore ∇_*p*_ is the gradient along ***p***_*h*_. Same as the bias field, we use the Gauss–Newton approach to update the initial velocity field *v*. Let

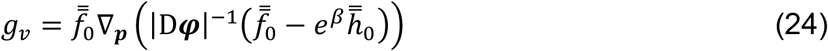

and

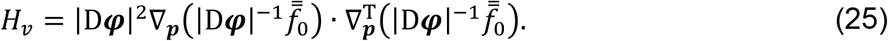

Also let ***b***_*v*_ = vec(*v*), ***g***_*v*_ = vec(*g_v_*), ***H***_*v*_ = diag(vec(*H_v_*)), the descent deviation is computed via

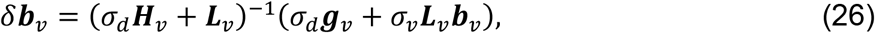

or in the 3D field form *δv* = ivec(*δ**b**_v_*), then the *v* field is updated by

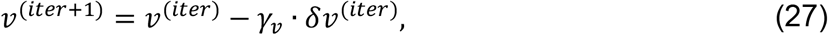

where 0 < *γ_v_* ≤ 1 controls the descent size. The matrix inversion of Eq. (26) is solved by the full multigrid algorithm (Ashburner, 2007). In this study, *L_v_* is defined with the same form as *L_β_*, where the strength of membrane energy and that of bending energy are controlled by *λ*_*v*,1_ and *λ*_*v*,2_, respectively. This differential operator is encoded in the matrix 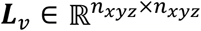.

#### 2.2.4 Optimization of *q_i_* and *c_i_*

Like the optimization of ***q***_0_, we use the Gauss-Newton approach (Press & Numerical Recipes, 2007) to update the values of ***q***_*i*_ and ***c***_*i*_. For each DW image *h_i_*, let

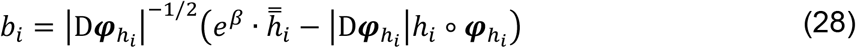

and ***b***_*i*_ = vec(*b_i_*). Also let

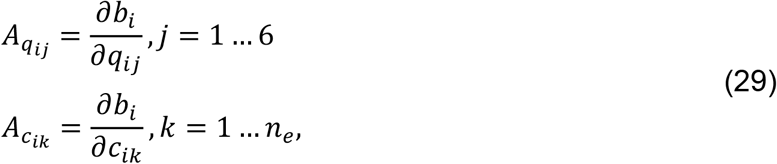

and 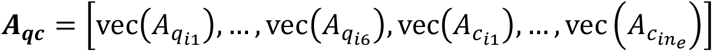. It follows that ***q***_*i*_ and ***c***_*t*_ are updated by

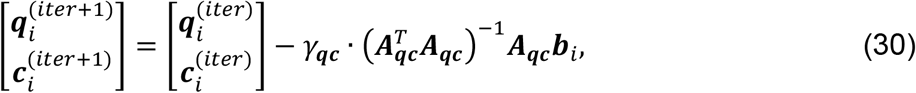

where 0 < *γ_qc_* ≤ 1.

### 2.3 Implementation

The proposed method is implemented with MATLAB R2020a (The MathWorks, Inc., Natick, Massachusetts, USA).

#### 2.3.1 Processing Anatomical Images

The anatomical image (T1w or T2w) is processed using the Segment toolbox of SPM12 (Ashburner & Friston, 2005). Briefly, this method uses a probabilistic framework to classify the input image into six components, including GM, WM, CSF, bone, scalp, and others, each is represented by a tissue probability map (TPM). A brain mask is constructed by combining the TPMs of GM, WM, and CSF. Note that the algorithm also conducts the correction for the intensity inhomogeneity, therefore, the processed T1w or T2w image is assumed to have negligible inhomogeneity.

#### 2.3.2 Estimating Diffusion Tensors and MAP-MRI Coefficients

The pseudo dMRI data is synthesized by either the Gaussian model of DTI (Basser et al., 1994) or the Hermite model of MAP-MRI (Özarslan et al., 2013). In the former, the diffusion tensors are estimated using the weighted linear least squares method (Koay et al., 2006). For the Hermite model, the diffusion signals are fitted by the linear version of the regularized MAP-MRI (ReMAP) algorithm (Hsu & Tseng, 2018). Although the linear methods may not achieve very good accuracy in tensor estimation (Jones & Cercignani, 2010) or in MAP-MRI estimation, they are sufficient for our purpose. First, our goal is not to precisely calculate the diffusion tensors or the Hermite coefficients, but to construct the pseudo DW images which have similar contrast to the raw DW images, in order for estimating the parameters of the EC-induced distortions and head motions. Second, the linear methods are very fast, therefore, synthesizing the pseudo dMRI data in each iteration would not become a computational burden.

#### 2.3.3 Solving 1D-LDDMM Problem

LDDMM is an elegant mathematical framework for the non-linear registration problem of large deformations, which is applicable to the susceptibility-induced distortions. Given the initial velocity, the large deformation could be constructed by solving Eq. (1), i.e., shooting the initial velocity from the identity at *t* = 0 to generate a series of velocity fields and associated deformations until *t* = 1. In practice, this is implemented by discretizing the time into *t_n_* uniform intervals (i.e., dividing the large deformation into *t_n_* small deformations), and integrating the velocities to create the deformations using, e.g., semi-Lagrangian method (Beg et al., 2005). Consequently, one of the drawbacks of the LDDMM framework is that the estimation demands a large amount of computation resources, which roughly increase with *t_n_*. In order to reduce the computational load, the current implementation sets *t_n_* to the minimum value, i.e., *t_n_* = 1. In this manner, the field map is simply 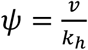. At first glance, it seems too simplified, however, in our experience, *t_n_* = 1 is sufficient to achieve good registration.

#### 2.3.4 Estimation Strategy

Since many parameters need to be estimated in the proposed integrated framework, we develop a multi-stage strategy to enhance the processing efficiency and reduce the chance of being trapped in local minima.

1. Stage 1. Fix *β* = 0 and *v* = 0, estimate the parameters for the head motions and EC-induced distortions by minimizing *E*_2_. This stage produces 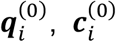, and 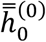.
2. Stage 2. Fix *β* = 0 and *v* = 0, conduct the rigid registration between 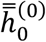 and the pseudo *b*_0_ image *f*_0_ by minimizing *E*_1_. This stage gives 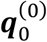.
3. Stage 3. Use 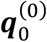, and *β* = 0, and *v* = 0 as the initial inputs for this stage. Conduct the registration between 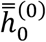 and *f*_0_ by minimizing *E*_1_. This stage produces 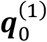, *β*^(0)^ and *v*^(0)^.
4. Stage 4. Fix *β* = *β*^(0)^ and *v* = *v*^(0)^, re-estimate the parameters for the head motions and EC-induced distortions by minimizing *E*_2_. This stage gives 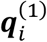 and 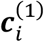.
5. Stage 5. Use 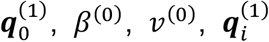, and 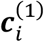 to initiate the iterative estimation in this stage; see Algorithm 1. In our experience, *E*_1_ or *E*_2_ in this stage does not always decrease monotonically during iterations even if the artefacts are reduced. Similar observation is reported in (Andersson & Sotiropoulos, 2016). Therefore, in this stage we set the iteration to a fixed number and ignore the values of *E*_1_ and *E*_2_ (Andersson & Sotiropoulos, 2016).

#### Algorithm 1

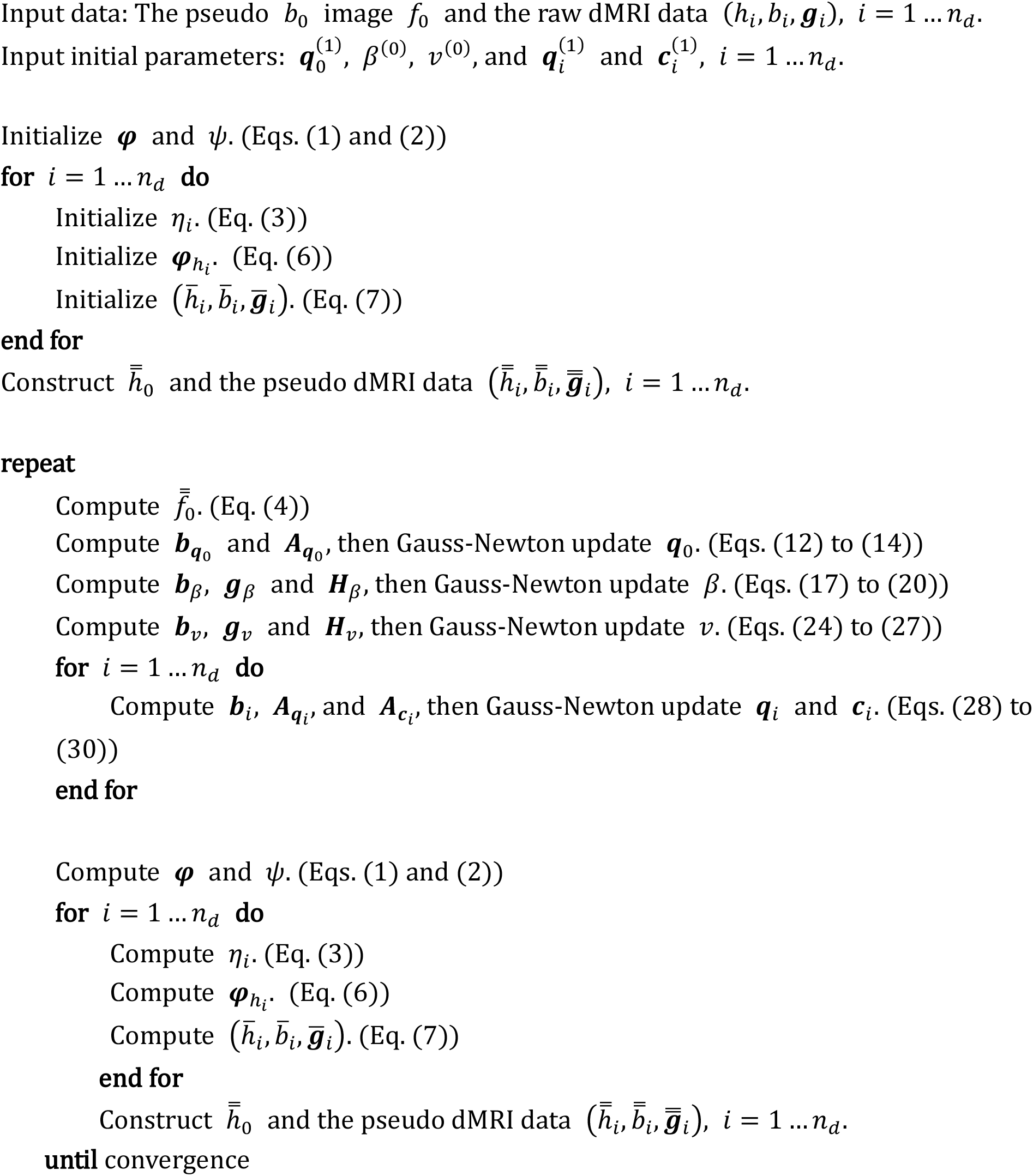

## 3. Evaluation

In the study, the performance of the DACO algorithm was assessed by two evaluations. The first one examined DACO using the real dMRI data from the human connectome project (HCP). Another evaluation used the real human dMRI data acquired from clinical scanners to validate the performance of DACO in the clinical settings. If not specified, the experiments performed in the evaluation used the same processing parameters. We used the quadratic model for the EC-induced magnetic fields.

### 3.1 Evaluation Using HCP Data

All of the HCP data were acquired on a Siemens Skyra 3T scanner (Siemens, Erlangen, Germany) which is specifically modified for the HCP project. The acquisition protocols of the T1w image, T2w image, and dMRI data were detailed elsewhere (Glasser et al., 2013; Sotiropoulos et al., 2013; Van Essen et al., 2012); here we briefly introduced the relevant information of the dMRI protocol. The single-shot spin-echo EPI sequence was used to acquire all images with parameters: TR/TE = 5520/89.5 ms, 111 axial slices with thickness = 1.25 mm, FOV = 210×180 mm^2^, in plane resolution = 1.25×1.25 mm^2^, echo spacing = 0.78 ms, partial Fourier = 6/8, and multiband factor = 3. A full dMRI data comprised two sets, one was acquired with the PE direction left to right (the LR set) and the other right to left (the RL set). The two sets used nearly identical monopolar diffusion-sensitive gradients, each comprised 18 *b*_0_ images and 3 shells of 90 DW images with b-values = 1000, 2000, and 3000 s/mm^2^, respectively. These images were intermixed and acquired in three sessions with 95, 96, and 97 images, respectively.

To ensure higher convenience and feasibility to use the HCP data, the HCP team officially released the T1w image, T2w image, and dMRI data with minimal preprocessing efforts (Glasser et al., 2013). Briefly, data of the six dMRI sessions (i.e., both the LR and RL sets) were concatenated and corrected using the HCP Pipeline, which comprised the TOPUP algorithm (Andersson et al., 2003) and the EDDY algorithm (Andersson & Sotiropoulos, 2016) implemented in FSL. The TOPUP algorithm was used to estimate the field map using the *b*_0_ images of the LR and RL sets, then the field map was input to the EDDY algorithm to correct the susceptibility-induced distortions, EC-induced distortions, and head motions. Besides, the HCP Pipeline merged the relevant DW images in the LR and RL sets to a single DW image to mitigate the pile-up problem (Andersson et al., 2003). Note that the official release of the artefact-corrected dMRI data and the T2w image had been registered to the T1w image.

The DACO algorithm was performed on the dMRI data of eight subjects. While the LR and RL sets were processed separately, the three dMRI sessions in each set were concatenated for artefact correction. Since TOPUP and EDDY are state-of-the-art algorithms for correcting the dMRI artefacts, we used the HCP Pipeline as the reference to evaluate the performance of the DACO algorithm. We used the normalized mutual information (NMI) (Studholme et al., 1999) over the entire brain to quantify the similarity between two images.

#### 3.1.1 Comparison between T1w and T2w

Since the pseudo *b*_0_ image plays an important role in the DACO algorithm, it is of interest to inquire which anatomical image, T1w or T2w, is better to synthesize the pseudo *b*_0_ image. We investigated this problem through calculating the NMI values between FA maps. Specifically, the DACO algorithm was conducted on the LR and RL sets using either the T1w or T2w image. For each subject, the FA maps of the LR set processed using the T1w image and the T2w image were produced, respectively. We calculated the NMI values of these FA maps with respect to the FA map derived from the dMRI data processed with the HCP Pipeline. The mean±standard deviation (SD) was 1.0669±0.0018 for the FA maps using T1w and 1.0664±0.0027 for the FA maps using T2w. The paired t-test on the NMI values showed that T1w- and T2w-based corrections were not statistically different (p-value=0.2354). The same calculation was performed on the RL set, the NMI values were 1.0681 ±0.0026 for T1w and 1.0683±0.0024 for T2w; both approaches were not statistically different (p-value=0.6566). The results revealed that, overall, using the T1w or T2w image to synthesize the pseudo *b*_0_ image achieved similar performance of artefact correction. In the study, we used the pseudo *b*_0_ image synthesized from the T1w image to conduct the following experiments.

#### 3.1.2 Evaluation of EC Parameters

The LR and RL sets of the HCP dMRI data used nearly the same diffusion encoding scheme, therefore, the EC-induced off-resonance magnetic fields of the relevant DW images in the two sets shall be close to each other. We made use of this property to evaluate the EC parameters by calculating the ten correlations, each for a EC parameter, between the LR and RL sets. We anticipated that the performance of the DACO algorithm was comparable to that of the HCP Pipeline because both methods used similar Gauss-Newton estimation approaches (Andersson & Sotiropoulos, 2016). Fig. 2 shows the results. We can see that the correlations of the EC parameters produced by DACO are high (>0.9) for those components not involving x (including y, z, y^2^, z^2^, and yz), moderate (>0.7) for the x^2^, xy, and xz components, and low (<0.3) for the x component and the constant term. The HCP Pipeline shows similar trends, but the discrepancies among the correlations are higher than the DACO algorithm: while components not involving x have higher correlations (>0.8), those involving x have substantially lower correlations (<0.3). The paired t-tests comparing every EC parameter between the DACO algorithm and the HCP Pipeline were conducted; the Benjamini-Hochberg method was used to control the false discovery rate (FDR) for all tests at a level of 0.05. Note that in the HCP Pipeline, the values of the constant term in the LR set were artificially set to be the same as those in the RL set, resulting in a correlation of one, therefore, the constant term was disregarded in the comparison. The comparison showed that the correlations of DACO were significantly higher than the counterparts of the HCP Pipeline. The HCP Pipeline fed both the LR and RL sets to the EDDY algorithm to conduct the artefact correction, whereas the DACO algorithm processed the two sets separately. The evaluation revealed that, even not sharing any information between the estimations for the LR and RL sets, the DACO algorithm still produced more consistent EC parameters than the HCP Pipeline, suggesting that the EC estimation of DACO was precise, which was comparable to, or even better than, that of the HCP Pipeline.

**Fig. 2.**
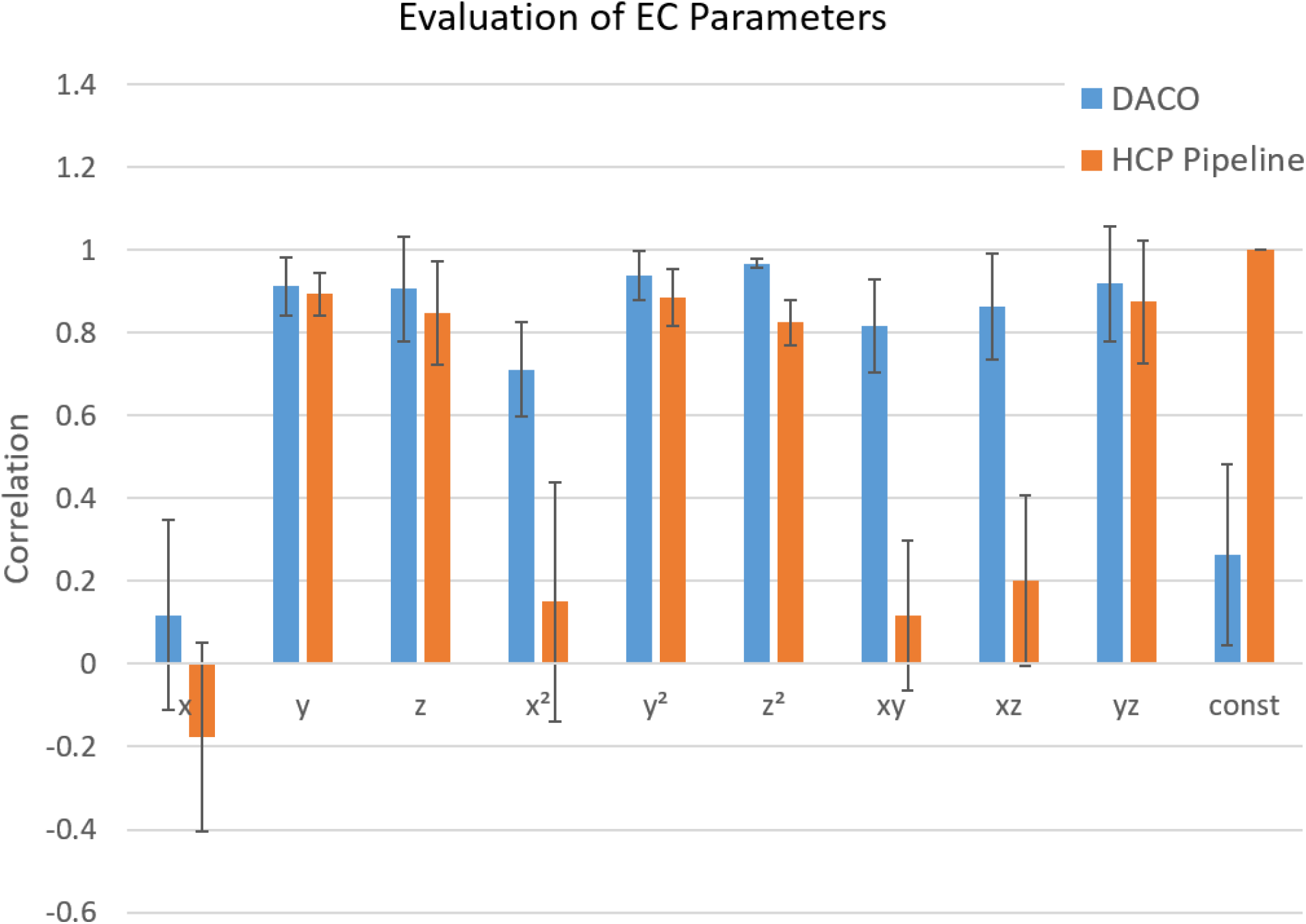
The correlations of the EC parameters between the LR and RL sets. The DACO algorithm has high correlations (>0.9) for those components not involving x (including y, z, y^2^, z^2^, and yz), moderate (>0.7) for the x^2^, xy, and xz components, and low (<0.3) for the x component and the constant term. The HCP Pipeline have higher correlations (>0.8) for components not involving x, and substantially lower correlations (<0.3) for those involving x. The correlations of DACO are significantly higher than the counterparts of the HCP Pipeline.

#### 3.1.3 Evaluation of Head Motion Parameters

In the evaluation, we calculated the six correlations (each for a head motion parameter) between the DACO algorithm and the HCP Pipeline. Since the two methods use slightly different definitions for the head motion parameters, for a proper comparison, the values of the HCP Pipeline have been transformed to the definition of DACO. Fig. 3 shows the comparison results for the LR and RL sets. The head motion parameters of DACO highly correlate (>0.8) with those of the HCP Pipeline, except for the x-translation component, where the correlations are moderate. The results suggest that the head motion parameters of DACO are similar to those of the HCP Pipeline in terms of the overall correlations.

**Fig. 3.**
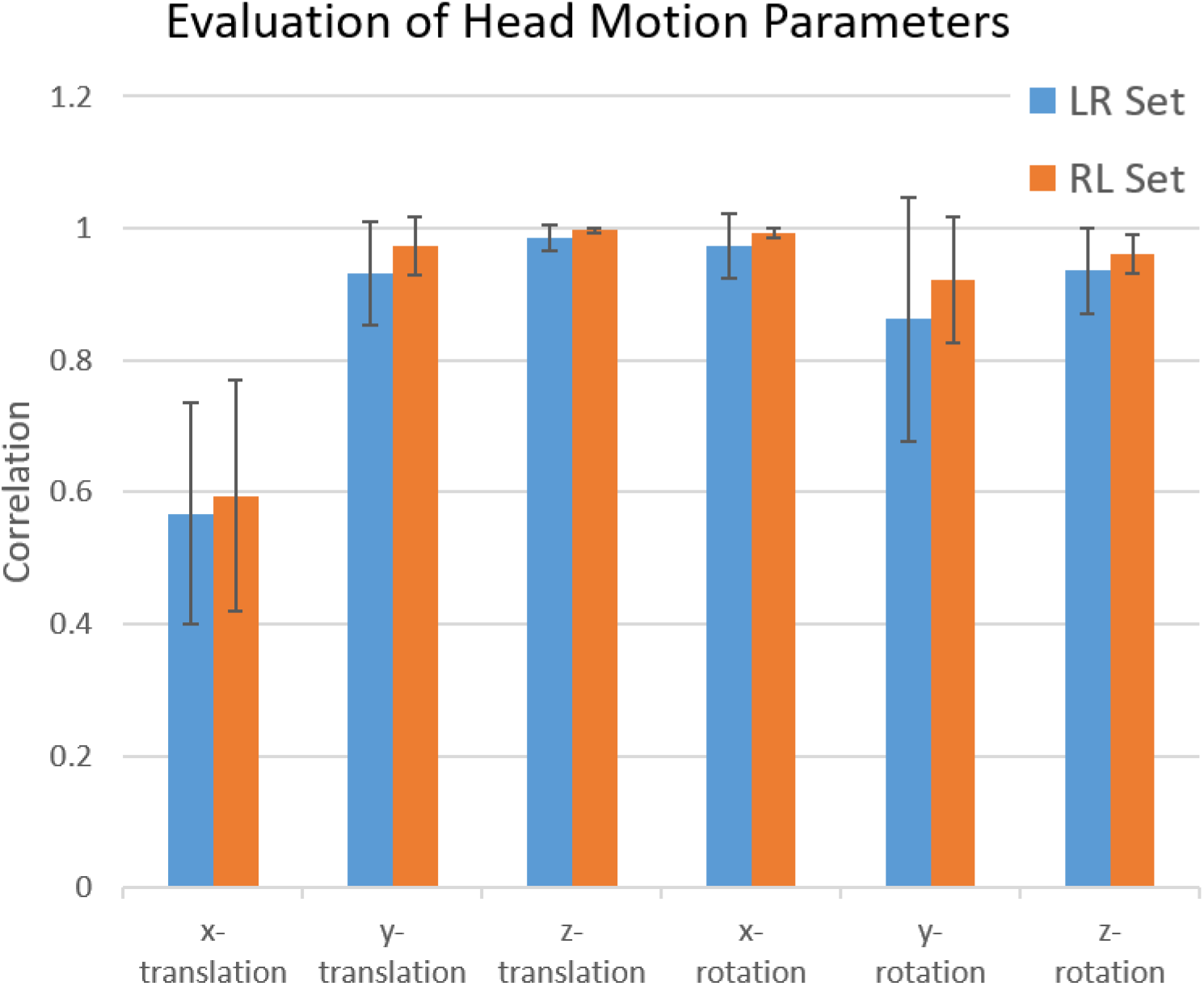
The correlations of head motion parameters between the DACO algorithm and the HCP Pipeline. The head motion parameters of DACO highly correlate (>0.8) with those of the HCP Pipeline, except for the x-translation component, where the correlations are moderate.

One of the approaches to study the performance of an algorithm in estimating the head motion parameters of dMRI data is to examine if the parameters vary smoothly (Andersson & Sotiropoulos, 2016). This approach assumes that the head moves slowly inside each scan session. The HCP dMRI data likely meet this assumption as this project recruited young adults who were cooperated to lay as still as possible during the scan sessions. Fig. 4 shows the head motion parameters of a typical subject produced by the DACO algorithm and the HCP Pipeline; the gray lines indicate the last images of scan sessions. Scrutinizing the time series of DACO, the gray lines exactly coincide the sudden “jumps” of the parameters, suggesting that the head motion parameters estimated by the DACO algorithm could adequately reflect the real scan conditions: while the head moves slowly inside the scan sessions, it may have abrupt movements between sessions. The HCP Pipeline has high correlations with the DACO algorithm, however, the HCP Pipeline has a subtle problem which is not revealed in the quantitative values. Specifically, we can observe that the HCP Pipeline has a series of equally distributed “spikes” along the time series in the y-rotation component (such as those indicated by the red arrows). All of the spikes coincide with the *b*_0_ images. This pattern contradicts to the assumption of slow motion inside the sessions; moreover, it is too regular to be true and could be considered as a sort of artefact. The results suggest that the DACO algorithm is slightly better than the HCP Pipeline in terms of the smoothness of the head motion parameters.

**Fig. 4.**
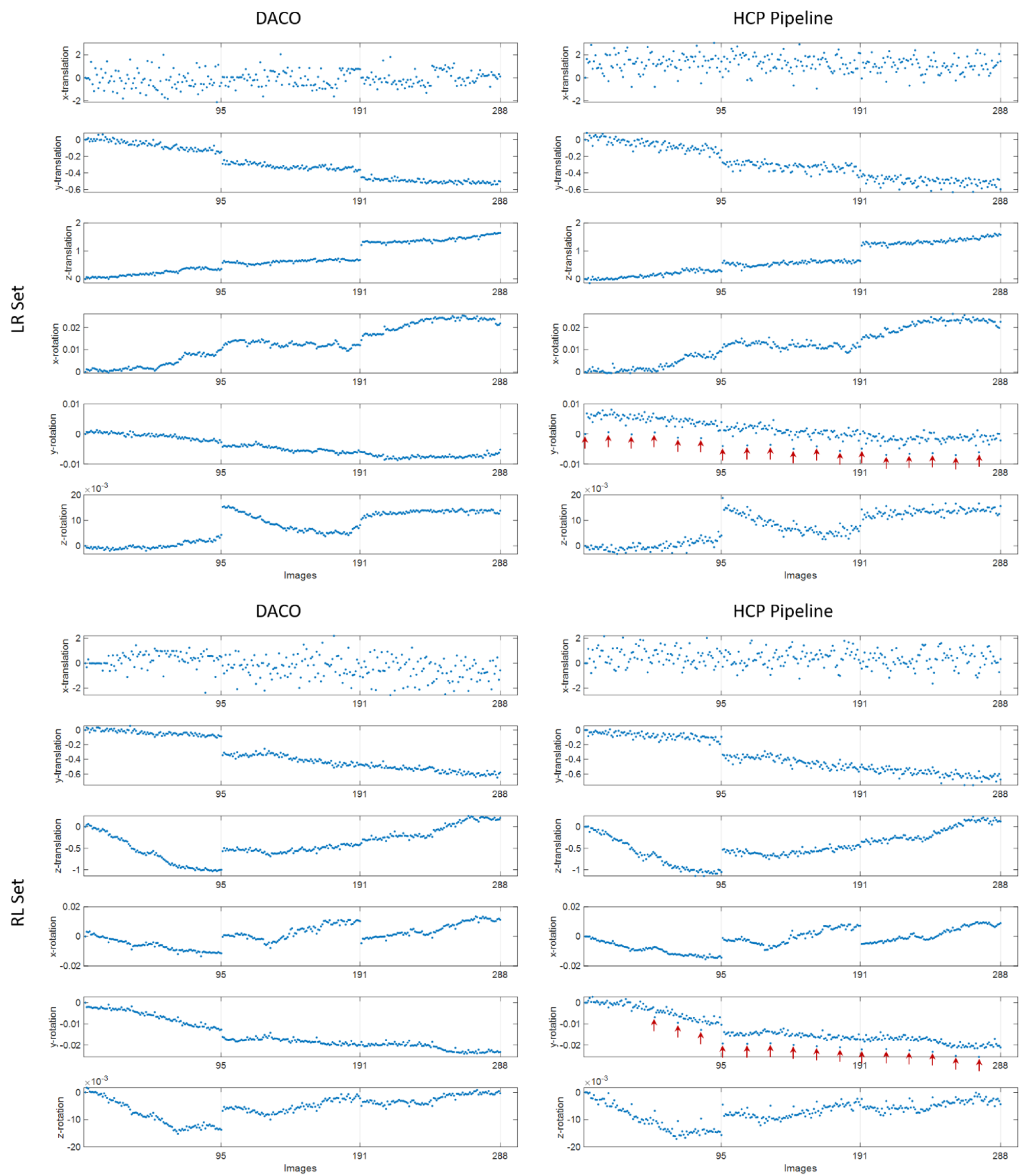
The head motion parameters of a typical subject of HCP. The left column is the DACO algorithm and right the HCP Pipeline. The gray lines indicate the last images of scan sessions. The gray lines exactly coincide the sudden “jumps” of the parameters of DACO. The HCP Pipeline has a series of equally distributed “spikes” along the time series in the y-rotation component (such as those indicated by the red arrows). All of the spikes coincide with the *b*_0_ images.

Among the head motion parameters, we can see that the x-translation component of DACO shows a nearly random pattern with large dispersion in both the LR and RL sets, which is greatly different from other components; the HCP Pipeline also exhibits similar patterns. This could be attributed to the fact that head motions would confound with the EC-induced distortions, particularly the translation along the PE direction ***p***_*h*_. Let the translation of head motions be decomposed into 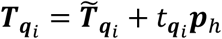, where *t_q_i__* is the component along ***p***_*h*_ and 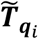 represents the residual translation. Rephrasing Eq. (6), the deformation of *h_i_* is expressed as

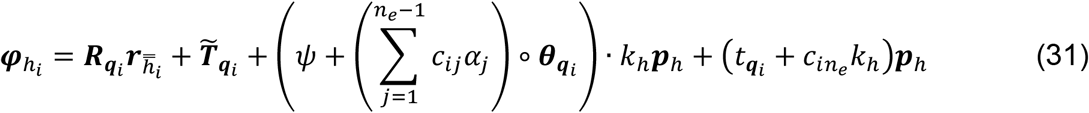

We can see that the rigid translation along the PE direction is represented by two parameters, one is from head motions (*t_q_i__*) and the other from the EC-induced distortions (*c_in_e__*, the constant term). Even though the eventual deformation ***φ***_*h_i_*_ could correct the dMRI artefacts properly, it is difficult to precisely determine the proportion contributed by each source, rendering their estimation not very accurate. In addition, the high variation of x-translation in the motion parameters partly explains the low correlations between the LR and RL sets in the EC parameters involving x and the constant term.

#### 3.1.4 Evaluation of Rigid Alignment between dMRI and T1w

In the DACO algorithm, we used SSD between the real and pseudo *b*_0_ images to quantify the rigid alignment between the space of dMRI and the space of T1w, where the ***q***_0_ at (locally) minimal SSD was considered as the best alignment. Previous study showed that SSD was more robust than NMI when the misalignment was large, however, NMI could achieve better alignment whenever it performed well (Bhushan et al., 2015). We employed this property to evaluate the rigid alignment of the DACO algorithm. Specifically, we rigidly aligned the FA map derived from the dMRI data which was corrected by the DACO algorithm to the WM TPM which was derived from the T1w image using SPM12 (Ashburner & Friston, 2005). We used NMI as the cost function to find the optimal alignment, where the optimization was conducted with the Powell’s method (Press & Numerical Recipes, 2007). In this manner, this optimal alignment was regarded as the misalignment of the DACO algorithm. Moreover, the magnitude of misalignment (in unit of mm) was calculated using the approach proposed by Power et. al. (Power et al., 2014). Similar procedures were conducted on the HCP Pipeline. The magnitudes of misalignment of DACO were 0.9638± 0.3248 mm and 0.6942±0.1631 mm for the LR set and RL set, respectively. The values of the HCP Pipeline were 0.9566±0.3119 mm. Paired t-tests showed that the DACO algorithm was not significantly different from the HCP Pipeline (p-values are 0.9735 and 0.0990 for the LR and RL sets, respectively). The results suggest that the DACO algorithm achieves similar performance to the HCP Pipeline in rigidly aligning the dMRI data to the anatomical image space.

#### 3.1.5 Evaluation of Bias Field

The evaluation of the bias field was conducted by calculating the similarity of the first *b*_0_ image of a dMRI data with respect to the pseudo *b*_0_ image, which is assumed to have negligible inhomogeneity artefact. Since the correction of intensity bias was a unique feature of the DACO algorithm, the HCP Pipeline was not employed as the reference. Instead, we calculated the NMI values of the raw *b*_0_ image (i.e., no correction applied), the *b*_0_ image of DACO with intensity bias retained (i.e., corrected with the field map), and the *b*_0_ image of DACO with intensity bias corrected (i.e., corrected with the field map and the bias field), with respect to the pseudo *b*_0_ image, respectively. The calculations were conducted for the LR and RL sets, respectively, and the results are shown in Fig. 5. The repeated analysis of variance (ANOVA) showed that the NMI values the LR and RL sets had significant differences (p-values<10^-5^). The *b*_0_ image of DACO with intensity bias corrected had the highest similarity to the pseudo *b*_0_ image, followed by the *b*_0_ image of DACO with intensity bias retained, and then the raw *b*_0_ image which had the lowest similarity. Fig. 6 shows the images of a typical subject; these images are respectively situated in the reference spaces of the LR set (upper row) and the RL set (bottom row). As we can see, while the raw *b*_0_ images (Fig. 6(b)(g)) suffer from significant susceptibility-induced distortions, they also reveal a pattern of higher intensity in the peripheral region and lower intensity in the central region. If we apply the field maps estimated by DACO, the distortions are substantially reduced, however, the intensity inhomogeneity is still prominent (Fig. 6(c)(h)). If we further apply the bias field, the intensity bias is substantially mitigated, rendering the appearance of the corrected *b*_0_ images (Fig. 6(d)(i)) highly similar to the pseudo *b*_0_ images (Fig. 6(a)(f)). In the DACO algorithm, the bias field is configured to correct both the intrinsic and the extrinsic intensity biases. Therefore, we can see that the estimated bias fields (Fig. 6(e)(j)) comprise not only a low frequency pattern, which is the characteristic of the extrinsic bias, but also a high frequency component, which is primarily due to the intrinsic differences between the real and pseudo *b*_0_ images. The results suggest that the bias field estimated by the DACO algorithm could adequately correct the intensity bias of dMRI data.

**Fig. 5.**
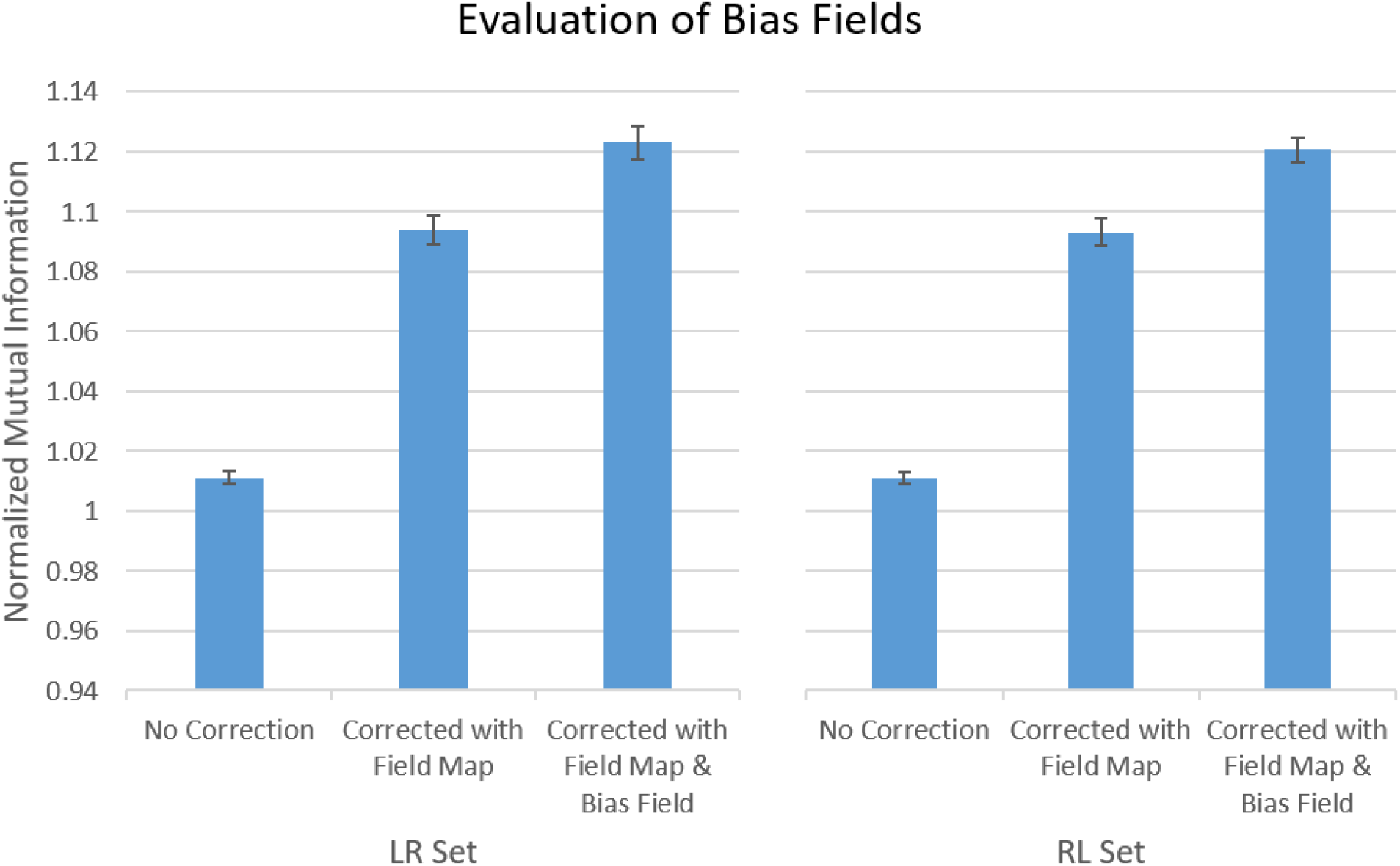
Quantitative evaluation for bias fields. The normalized mutual information (NMI) was used to measure the similarity between the first *b*_0_ image of a dMRI data with respect to the pseudo *b*_0_ image, which is assumed to have negligible inhomogeneity artefact. The measurement was conducted on the raw *b*_0_ image (No Correction), the *b*_0_ image of DACO with intensity biases retained (Field Map), and the *b*_0_ image of DACO with intensity biases corrected (Field Map & Bias Field).

**Fig. 6.**
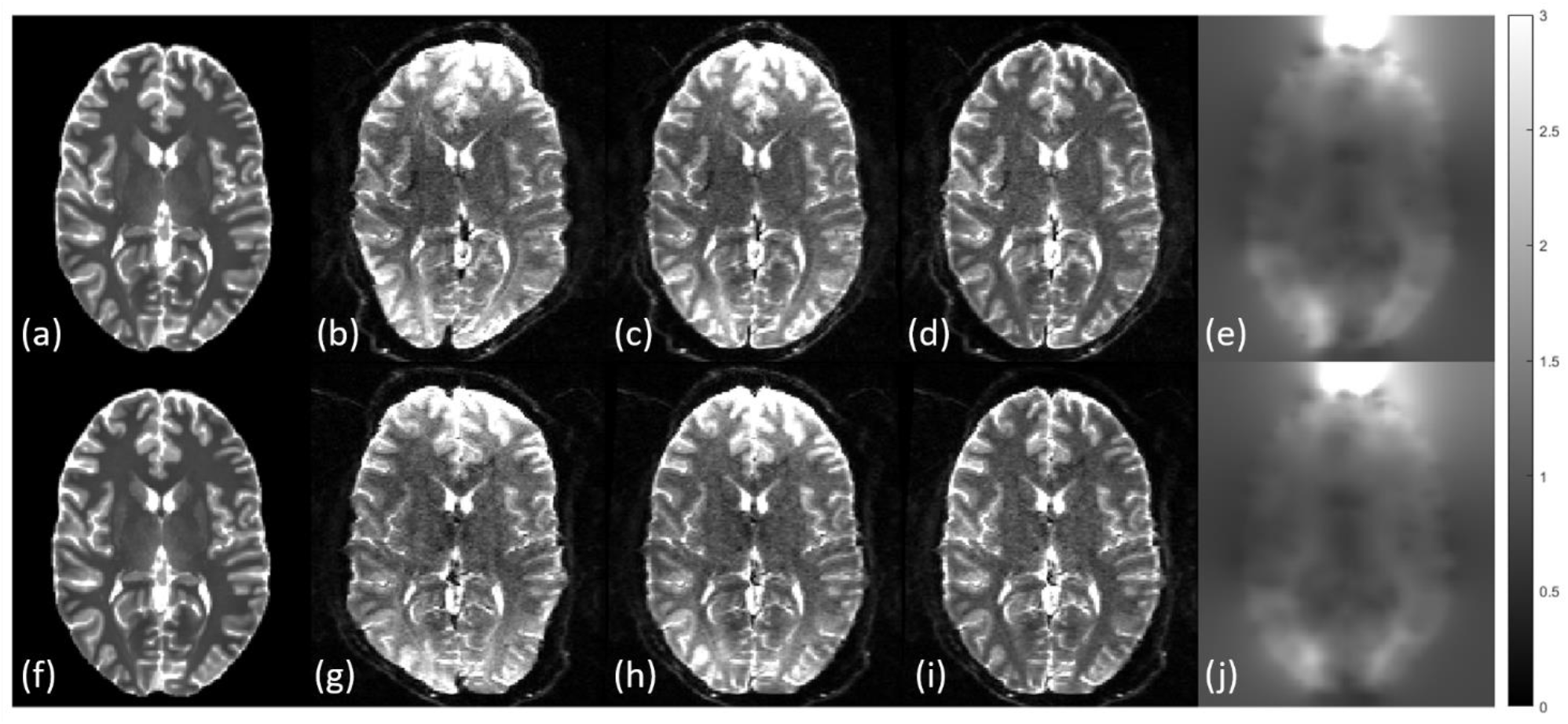
Qualitative assessment for bias fields with a typical subject of HCP. The images are situated in the reference spaces of the LR set (upper row) and the RL set (bottom row), respectively, with (a)(f) the pseudo *b*_0_ images, (b)(g) the raw *b*_0_ images, (c)(h) the *b*_0_images corrected with the field maps of DACO, (d)(i) the *b*_0_ images corrected with the field maps and bias fields of DACO, and (e)(j) the bias fields estimated by DACO.

#### 3.1.6 Evaluation of Field Map

The performance of DACO in correcting the susceptibility-induced distortions is evaluated by measuring the similarity of the first *b*_0_ image of a dMRI data with reference to the T2w image, of which the intensity inhomogeneity was retained and the non-brain regions were masked out. We calculated the NMI values of the raw *b*_0_ image (i.e., no correction applied), the *b*_0_ image of DACO with intensity biases retained (i.e., corrected with the field map of DACO), and the *b*_0_ image of HCP Pipeline (i.e., corrected with the field map of HCP Pipeline), with respect to the T2w image, respectively. The results are shown in Fig. 7. In either the LR or RL set, the repeated ANOVA shows that the NMI values among the *b*_0_images of no correction, DACO, and HCP Pipeline have significant differences (both p-values<10^-5^). The *b*_0_ image of HCP Pipeline has the highest NMI value (1.1163±0.0047), followed by the *b*_0_ images of DACO with the intensity biases retained (1.0878±0.0033 for LR and 1.0853±0.0040 for RL), and least the raw *b*_0_ image (1.0126±0.0022 for LR and 1.0123±0.0019 for RL). In terms of the similarity with reference to the T2w image measured by the NMI values, the DACO algorithm achieves 97.44±0.26% of the HCP Pipeline for the LR set and 97.22±0.44% for the RL set. Using the NMI values of the raw *b*_0_ images as the baseline, the improvement of geometrical consistency of the DACO algorithm is 72.50± 2.43% of that of the HCP Pipeline for the LR set, and 70.20±4.46% for the RL set.

**Fig. 7.**
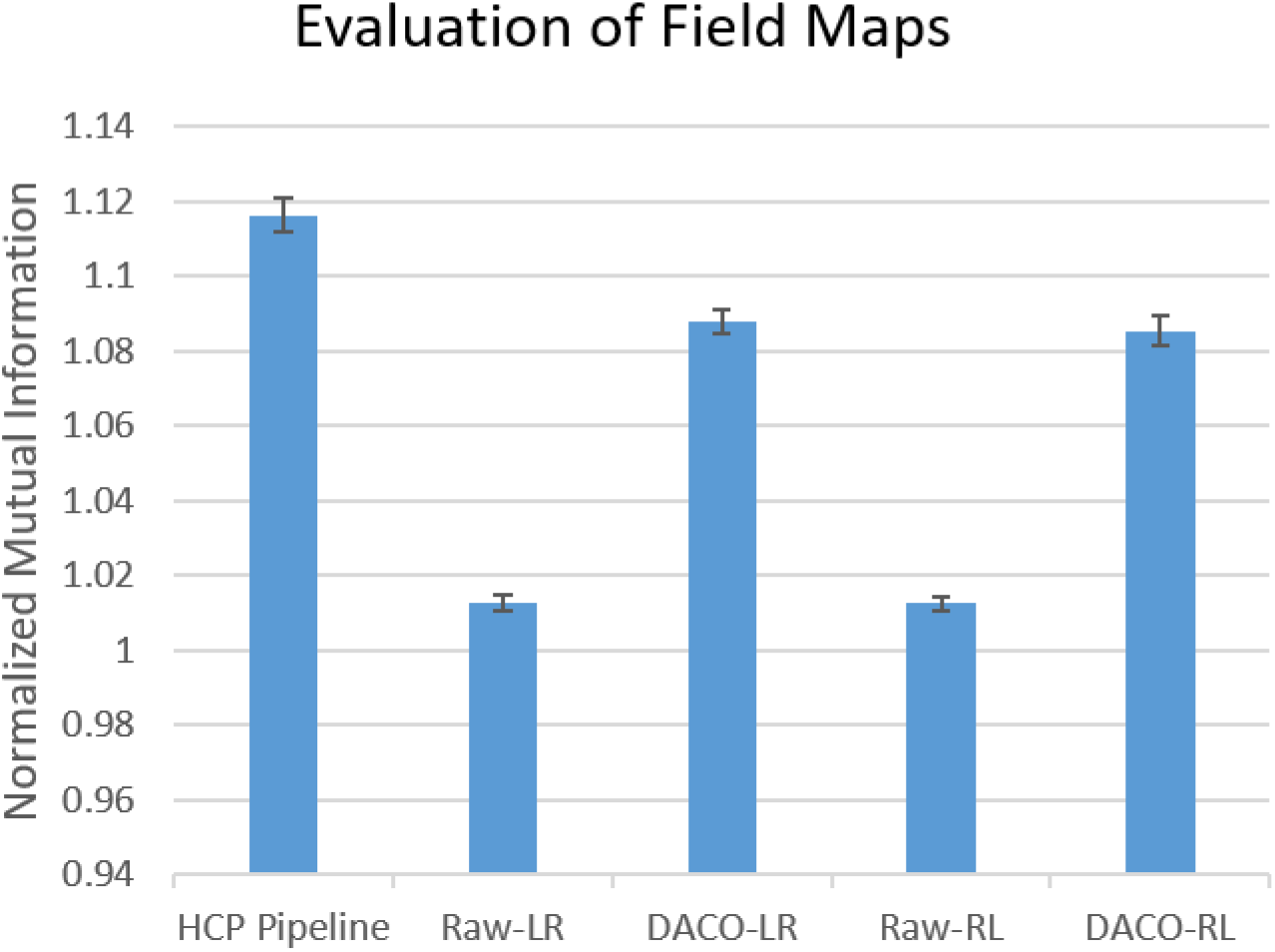
Quantitative evaluation for field maps. The normalized mutual information (NMI) was used to measure the similarity between the first *b*_0_ image of a dMRI data with reference to the T2w image, of which the intensity inhomogeneity was retained and the non-brain regions were masked out. The measurement was conducted on the raw *b*_0_ images of the LR set and RL set (Raw-LR and Raw-RL, respectively), the *b*_0_ images of DACO with intensity biases retained for the LR set and RL set (DACO-LR and DACO-RL, respectively), and the *b*_0_ image of the HCP Pipeline (i.e., corrected with the field map of HCP Pipeline).

Fig. 8 shows the images of a typical subject; these images are respectively situated in the reference spaces of the LR set (upper row) and the RL set (bottom row). We can see that the raw *b*_0_ images (Fig. 8(b)(g)) are corrupted by severe susceptibility-induced distortions in the left-right direction. The distortions are substantially mitigated by the DACO algorithm and the HCP Pipeline, so that the *b*_0_ images of DACO with the intensity bias retained (Fig. 8(c)(h)) and the *b*_0_ images of HCP Pipeline (Fig. 8(d)(i)) are very similar to the T2w images (Fig. 8(a)(f)). However, we can observe that the *b*_0_ images of HCP Pipeline exhibits higher similarity than the *b*_0_ images of DACO with the intensity bias retained, consistent with the quantification results. The field map of the LR set (Fig. 8(e)) and that of the RL set (Fig. 8(j)) are generally similar, but in regions with high susceptibility-induced off-resonance magnetic fields (such as those indicated by red circles) the two maps have higher discrepancies.

**Fig. 8.**
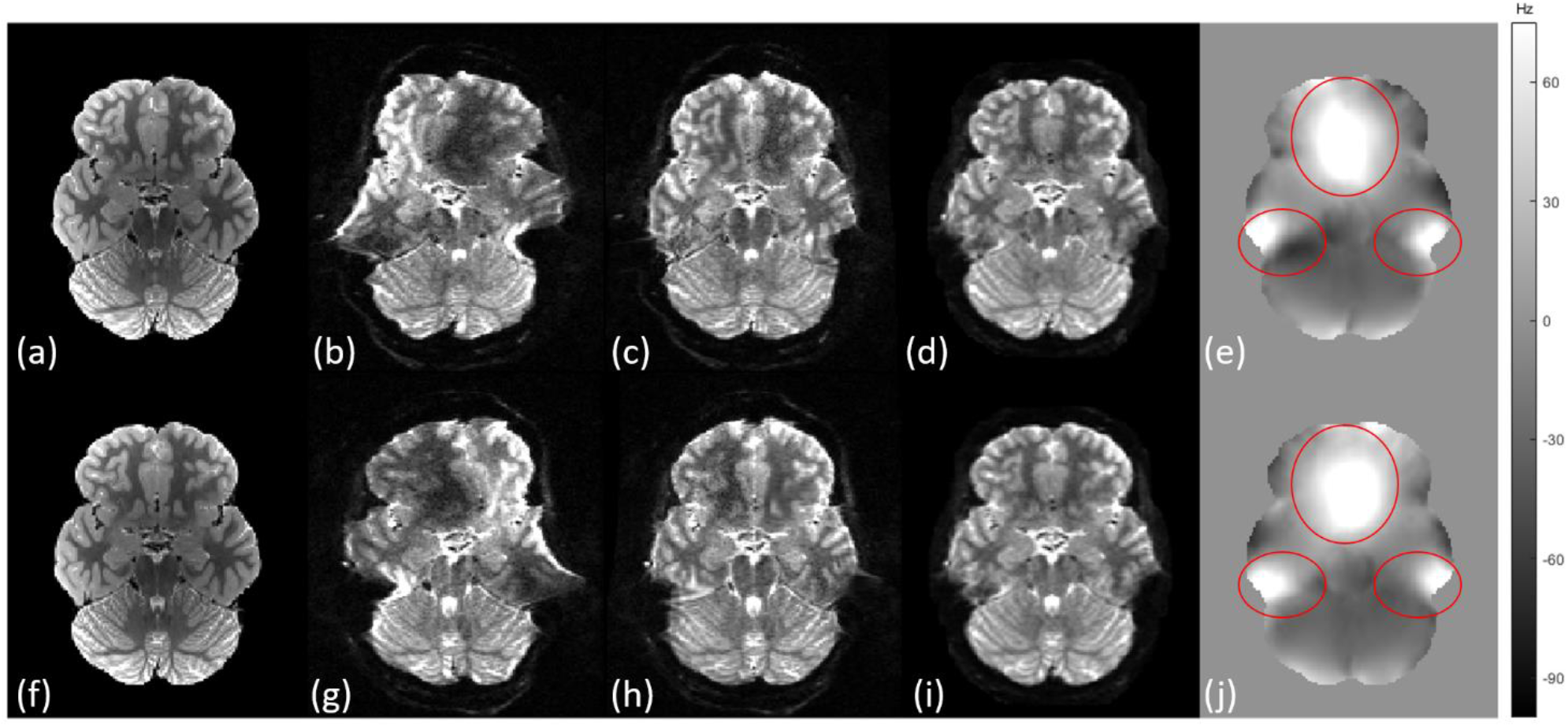
Qualitative assessment for field maps with a typical subject of HCP. The images are situated on the reference spaces of the LR set (upper row) and the RL set (bottom row), respectively, with (a)(f) the pseudo *b*_0_ images, (b)(g) the raw *b*_0_ images, (c)(h) the *b*_0_images corrected with the field maps of DACO, (d)(i) the *b*_0_ images corrected with the field map of the HCP Pipeline, and (e)(j) the field maps estimated by DACO. Red circles indicate those regions with higher discrepancies between the field maps of the LR set and the RL set.

The evaluation reveals that the DACO algorithm could effectively correct the susceptibility-induced distortions, but it does not perform as well as the HCP Pipeline. This is reasonable because the TOPUP algorithm of the HCP Pipeline estimates the field maps based on a strong physical constraint that the *b*_0_ images with two opposite PE directions have distortion patterns opposite to each other. In contrast, the DACO algorithm estimates the field maps based on the similarity between the real and pseudo *b*_0_ images, which is a much weaker constraint. Another reason why the DACO correction is less satisfactory is that the HCP Pipeline combines both the LR and RL sets to mitigate the pile-up problem (Andersson et al., 2003), therefore, the intensity of the *b*_0_ image (and other DW images) is properly restored, making the image greatly similar to the T2w image. In contrast, the DACO algorithm processes the LR and RL sets separately, and so it cannot restore the intensity adequately as does the HCP Pipeline, particularly in regions with severe pile-up problems.

#### 3.2 Evaluation Using Human Brain Data Acquired from Clinical Scanners

This evaluation used three dMRI data acquired from clinical scanners of different manufactures to examine the feasibility of DACO in clinical settings. Dataset A was selected from the IXI-HH cohort (*IXI dataset:* https://brain-development.org/ixi-dataset/) which used a Philips Intera 3T scanner (Philips Healthcare, Best, The Netherlands). Dataset B was a healthy subject from the OASIS-3 database (LaMontagne et al., 2019), where the MRI scans were collected on a Siemens TIM Trio 3T scanner with a 20-channel head coil. Dataset C I was acquired on a GE Discovery MR750w 3T scanner (GE Medical System, USA) with a 24-channel head coil. The anatomical images used in the evaluation were all T1w images. The acquisition parameters of T1w images and dMRI data were listed in Table 1. To summarize the performance of artefact correction, we compared the FA maps of the dMRI data to the WM TPM, which was derived from the T1w image using the Segment toolbox ofSPM12 (Ashburner & Friston, 2005). Since WM TPM and FA depicted the WM regions (Hsu et al., 2015) and had similar appearance, the WM TPM was used as the surrogate of FA without distortions.

**Table 1.** Acquisition parameters of the clinical data.

|  |  | Dataset A | Dataset B | Dataset C |
| --- | --- | --- | --- | --- |
| Scanner | Manufacturer | Philips | Siemens | GE |
|  | Model | Intera | TIM Trio | Discovery MR750w |
|  | Strength (Tesla) | 3 | 3 | 3 |
| Diffusion | TR/TE (ms) | 11894.4/51 | 14500/112 | 11500/100 |
|  | Voxel size (mm<sup>3</sup>) | 1.75×1.75×2 | 2×2×2 | 2×2×2 |
|  | Effective echo spacing (ms) | 0.228* | 0.980 | 0.744 |
|  | Phase-encoding direction | PA* | AP | PA |
|  | # diffusion directions | 16 | 24** | 61 |
|  | b-max (s/mm<sup>2</sup>) | 1000 | 1400 | 1000 |
| T1w | TR/TE/TI (ms) | 9.6/4.6/NA | 2400/1000/3.16 | 8.5/3.2/450 |
|  | Flip angle (°) | 8 | 8 | 12 |
|  | Voxel size (mm<sup>3</sup>) | 1×1×1 | 1×1×1 | 1×1×1 |

PA: posterior to anterior; AP: anterior to posterior.
\* The IXI does not officially provide the phase-encoding direction and the effective echo spacing, here we assume the former is PA and the latter is 0.228 ms.
\*\* Two dMRI sessions were conducted with exactly the same scanning parameters.

Fig. 9 shows the results of Dataset A. The pseudo *b*_0_ image (Fig. 9(e)), constructed from the T1w image (Fig. 9(a)), has similar appearance to the real *b*_0_ image (Fig. 9(f)). While the raw *b*_0_ image (Fig. 9(b)) shows strong susceptibility-induced distortions, the raw DW images (Fig. 9(c) and (d)) are additionally corrupted with the EC-induced distortions, rendering them respectively show a compression and an elongation along the PE direction (anterior-posterior). The bias field (Fig. 9(i)), the filed map (Fig. 9(j)), and the EC-induced fields (Fig. 9(k) and (l)) were estimated by the DACO algorithm. Applying these models to Dataset A, we can observe that the artefacts are substantially mitigated, rendering the *b*_0_image (Fig. 9(f)) and the DW images (Fig. 9(g) and (h)) highly matched to the red brain contour drawn from the T1w image. The artefact reduction is also evident from the FA maps (Fig. 9(n), (o), and (p)). A rim with elevated FA values is a common phenomenon usually observed in dMRI data suffering from significant EC-induced distortions and/or head motions (Jones & Cercignani, 2010). We can see that the FA map with correction only on head motions (Fig. 9(n)) has the most severe artefact. The rim is slightly reduced in the FA map with correction on the head motions and EC-induced distortions (Fig. 9(o)), however the rim is still prominent. The rim nearly disappears in the FA map with correction on the head motion, EC-induced distortions, and susceptibility-induced distortions (i.e., the DACO algorithm, Fig. 9(p)). In addition to the rim, using the red WM contours drawn from the WM TPM (Fig. 9(m)) as the reference, we can see that the WM contour outlined in the FA map of DACO (Fig. 9(p)) aligns with the WM TPM contour very well. In contrast, the contours in other FA maps (Fig. 9(n) and (o)) are not well aligned.

**Fig. 9.**
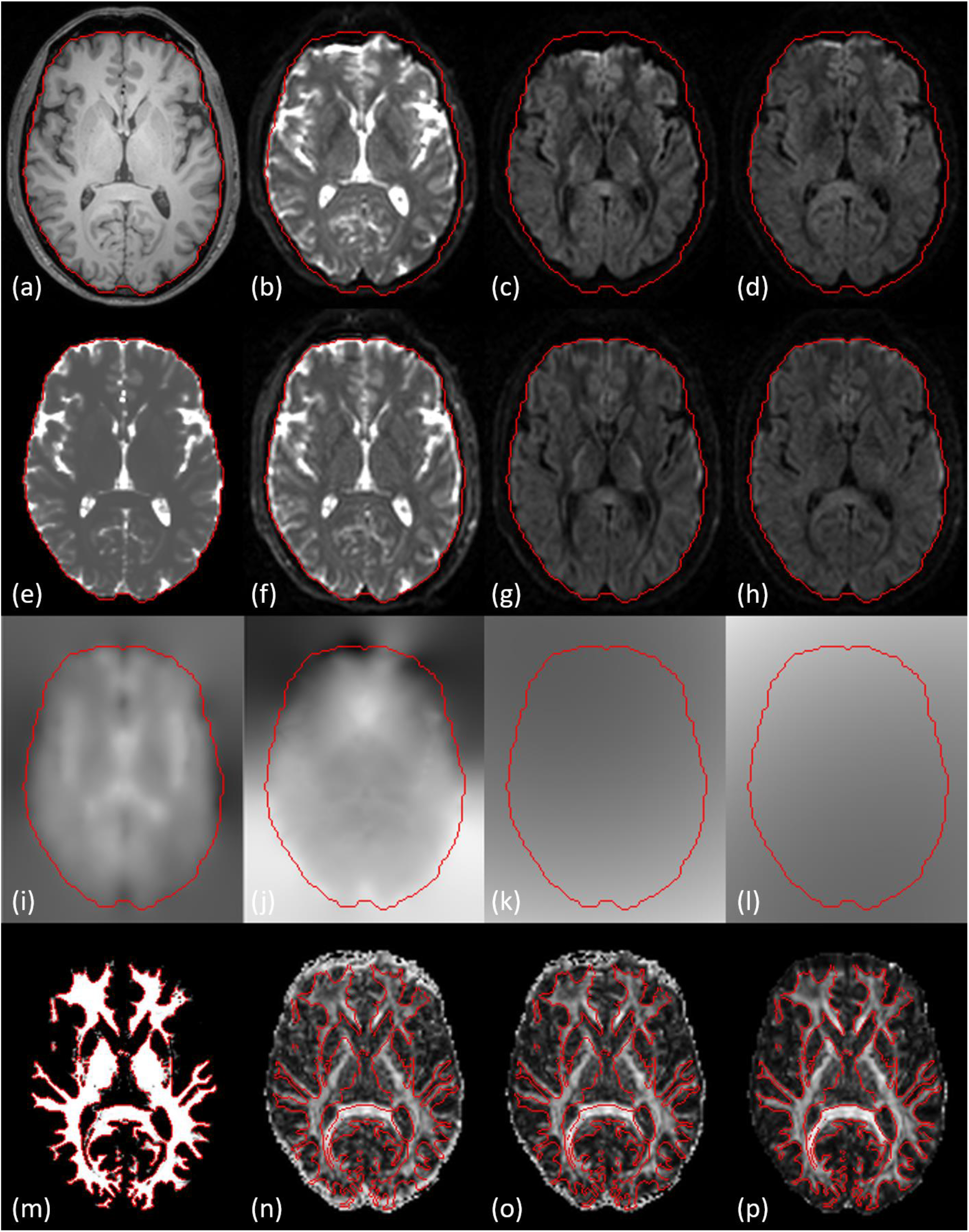
Processing results of Dataset A. The images are (a) the T1w image, (b) the raw *b*_0_image, (c)(d) the raw DW images, (e) the pseudo *b*_0_ image, (f) the corrected *b*_0_ image, (g)(h) the corrected DW images, (i) the bias field, (j) the filed map, (k)(l) the EC-induced fields, (m) the tissue probability map of white matter, (n) the FA map with correction only on head motions, (o) the FA map with correction on the head motions and EC-induced distortions, (p) the FA map with correction on the head motions, EC-induced distortions, and susceptibility-induced distortions (i.e., the DACO algorithm). The brain contours of (a) to (l) are drawn from (e). The white matter contours of (m) to (p) are drawn from (m).

One of the features of DACO is that we developed a dedicated registration algorithm for the susceptibility-induced distortions, which uses a Gauss-Newton optimization scheme (Eqs. (24) to (27)). To assess its effectiveness and efficiency, we compared this algorithm with another optimization scheme, the gradient descent (GD). Specifically, we replaced Eq. (26) with

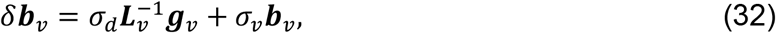

where ***g***_*v*_ was computed by Eq. (24), and kept other settings unchanged, and referred this algorithm to DACO-GD. We applied the DACO and DACO-GD algorithms to Dataset B. As shown in Fig. 10, both DACO (top row) and DACO-GD (middle row) substantially mitigate the large susceptibility-induced distortions observed in the raw *b*_0_ image (Fig. 10(a)). In addition, the algorithms estimate similar bias fields, field maps, and parameters for the head motions, EC-induced distortions, and rigid alignment. Therefore, the aligned pseudo *b*_0_images (Fig. 10(b) and (g)), the corrected *b*_0_ images (Fig. 10(c) and (h)), the FA maps (Fig. 10(d) and (i)), the bias fields (Fig. 10(e) and (j)), and the field maps (Fig. 10(f) and (k)) have small absolute differences (Fig. 10(l) to (p), respectively). Table 2 shows the processing time of the two algorithms in each stage. Although they had very similar performance in distortion correction, they were substantially different in the processing time of the Stage 3. While DACO spent 20.60 min to converge, DACO-GD spent 132.30 min, which was more than 6 folds to DACO. Our results show that the Gauss-Newton optimization scheme proposed in the study is an efficient and effective approach to estimate the field map. The results also reveal that the hessian equation (Eq. (25)) works well in the Gauss-Newton optimization scheme, even though the hessian is not derived rigorously but through an empirical approximation (see Appendix A).

**Fig. 10.**
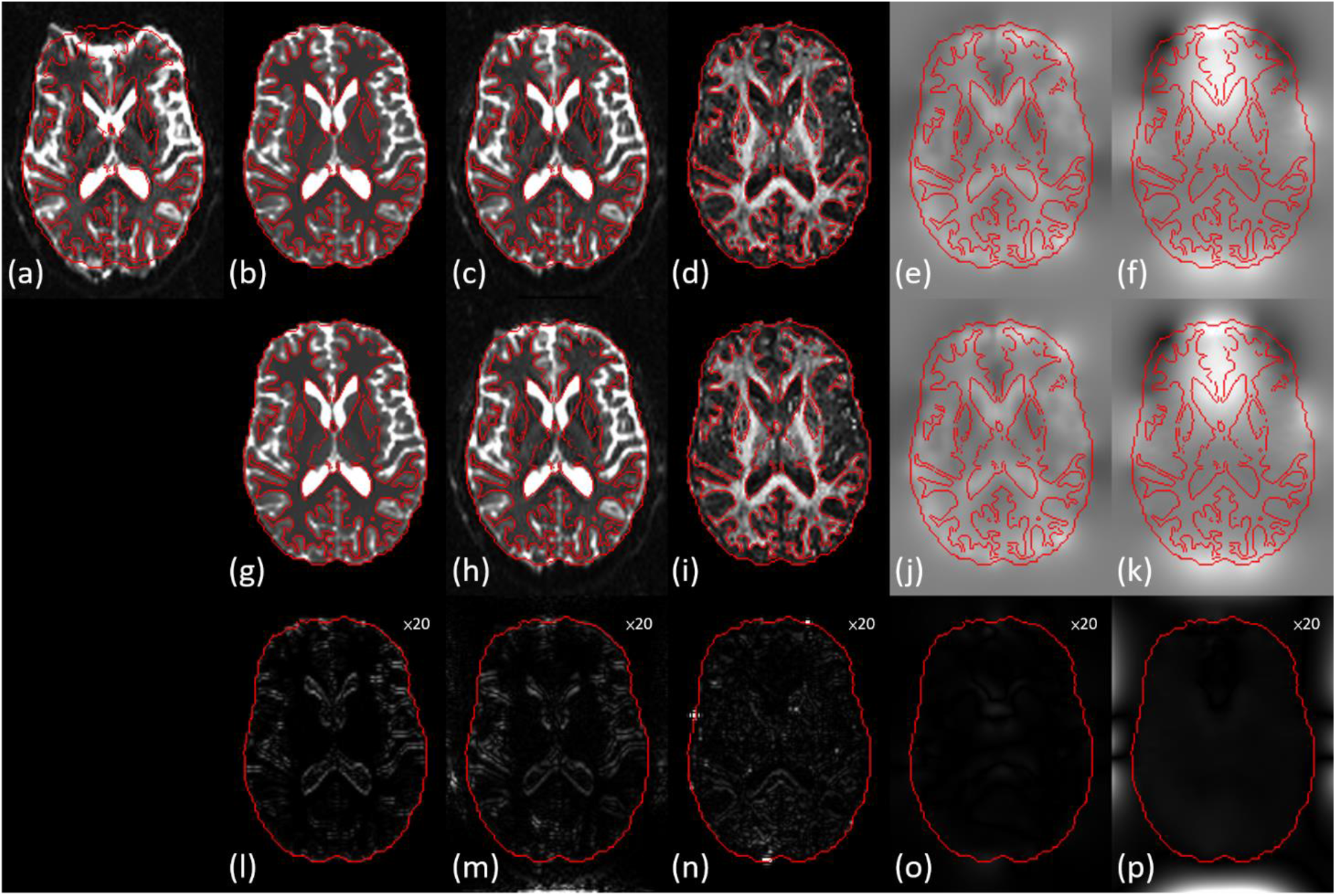
Processing results of Dataset B. Besides (a) the raw *b*_0_ image, the top and middle rows are the results of DACO and DACO-GD algorithms, respectively, with (b)(g) the aligned pseudo *b*_0_ images, (c)(h) the corrected *b*_0_ images, (d)(i) the FA maps, (e)(j) the bias fields, and (f)(k) the field maps. The bottom row are the absolute differences between DACO and DACO-GD, with the values magnified 20 folds for ease of visualization. The brain contours are drawn from (b) and the white matter contours are drawn from the tissue probability map of white matter.

**Table 2.** Processing time of each stage (in unit of minute).

|  | Stage 1 | Stage 2 | Stage 3 | Stage 4 | Stage 5 | Total |
| --- | --- | --- | --- | --- | --- | --- |
| DACO | 3.10 | 0.73 | 20.60 | 1.73 | 14.30 | 40.46 |
| DACO-GD | 3.08 | 0.73 | 132.30 | 1.72 | 13.88 | 151.71 |

Fig. 11 shows the results of Dataset C. Previous experiment shows that considering the intensity bias in the artefact models can significantly mitigate the intensity inhomogeneity. Also, modeling the intensity bias may lower the chance of being trapped in local minima and subsequently facilitate a good correction on the susceptibility-induced distortions. Fig. 11 demonstrates an example. The raw *b*_0_ image (Fig. 11(a)) suffers from large distortions around the frontal regions, with reference to the pseudo *b*_0_ image (Fig. 11 (e)). If the artefact model does not take the intensity bias into consideration, the algorithm falls into local minima, rendering the distortions not adequately corrected, such as those regions indicated by the green arrows in the *b*_0_ image (Fig. 11(b)) and the FA map (Fig. 11(c)). On the other hand, using the bias field (Fig. 11 (i)) to model the intensity biases can result in a proper registration, hence the *b*_0_ image (Fig. 11(f)) and the FA map (Fig. 11(g)) aligned to the pseudo *b*_0_ image very well. The field map estimated using the model with intensity biases (Fig. 11(h)) is much smoother than that without (Fig. 11(d)). An explanation to the results is the locally large image discrepancy between the real and pseudo *b*_0_ images. Recall that three primary sources contribute to the image discrepancy: rigid misalignment (by rigid head movement), spatial discrepancy (by the susceptibility-induced distortions), and the contrast discrepancy (by intensity bias). If the algorithm disregards the contribution of intensity bias, the algorithm would attempt to reduce the cost function (i.e., SSD) through spatially aligning the registered images. It is well known that registration, particularly non-linear, is an ill-posed problem. It follows that registering regions with large intensity bias may easily result in wrong point-by-point correspondence and wrong field map. This example demonstrates that modeling the intensity bias is helpful for correcting the susceptibility-induced distortions.

**Fig. 11.**
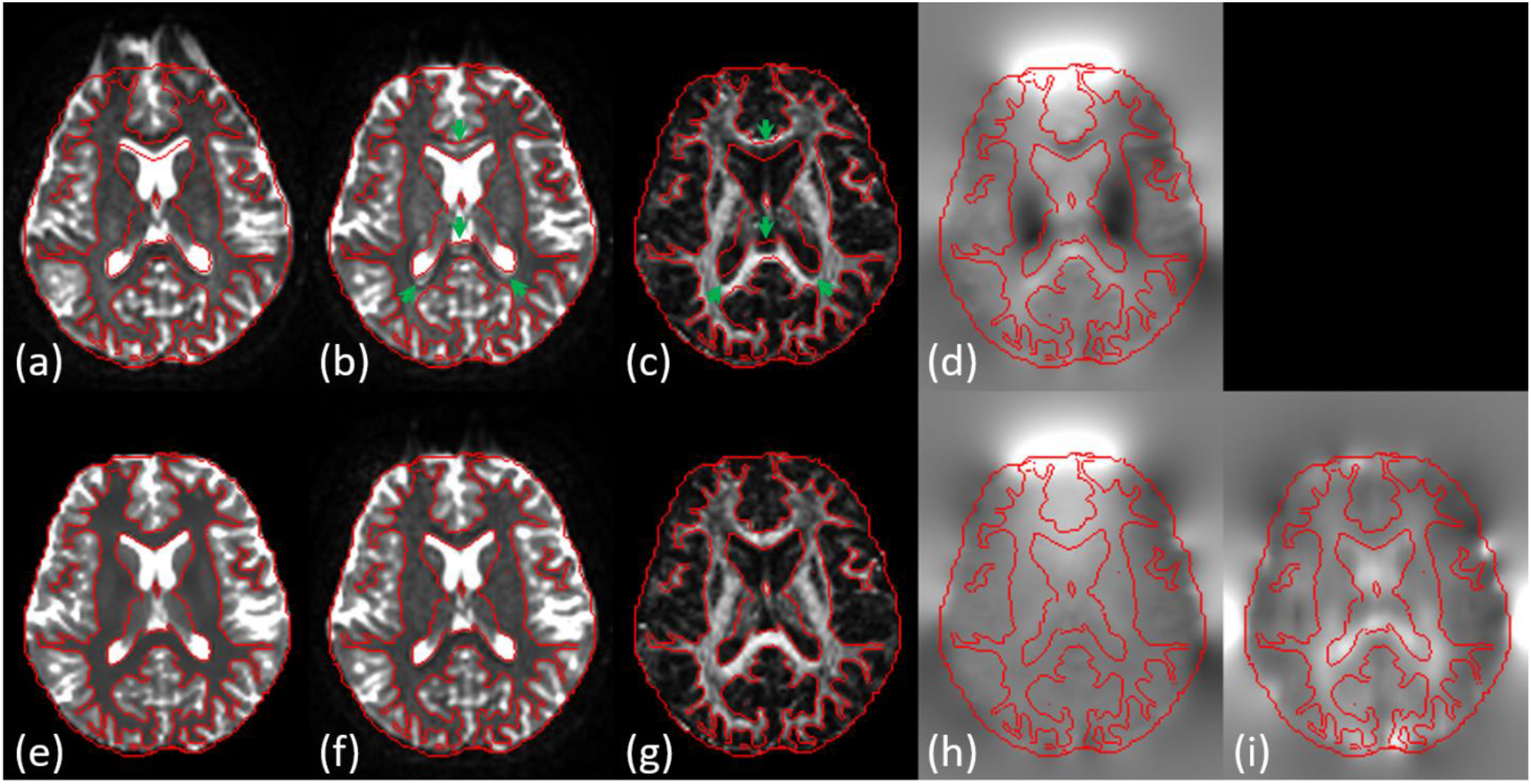
Processing results of Dataset C. The top row shows (a) the raw *b*_0_ image and the results of not considering the intensity bias, including (b) the corrected *b*_0_ image, (c) the FA map, and (d) the field map. The bottom row shows (e) the pseudo *b*_0_ image and the results of considering the intensity biases, including (f) the corrected *b*_0_ image, (g) the FA map, (h) the field map, and (i) the bias field. The green arrows indicate the regions where the susceptibility-induced distortions not adequately corrected. The brain contours are drawn from (e) and the white matter contours are drawn from the tissue probability map of white matter.

### 4. Discussion

In this study, we developed and implemented the DACO algorithm to correct the notorious distortions/artefacts of dMRI data, including the susceptibility-induced distortions, EC-induced distortions, head motions, intensity inhomogeneity, and the rigid misalignment between the dMRI data and anatomical image. Performance of this algorithm was evaluated on the HCP data. The results showed that DACO was comparable to the HCP Pipeline in correcting the EC-induced distortions, head motions, and rigid misalignment. The evaluation also revealed that DACO could substantially mitigate the susceptibility-induced distortions, even though its performance was slightly inferior to the HCP Pipeline. For the intensity inhomogeneity, the correction of DACO was adequate as well. The algorithm was also applied on three real human data acquired from clinical scanners of different manufactures, the results showed that this method properly corrected these artefacts, revealing its potential in clinical applications.

### 4.1 Contributions

Our method has three significant contributions. First, while previous algorithms have been proposed to address one or some dMRI artefacts, to the best of our knowledge, DACO is the first to address these artefacts altogether in an integrated framework. The necessity of addressing the artefacts altogether is that these artefacts interact with each other. For example, the susceptibility-induced distortions suffered by each of the DW images would be slightly different because of the head movements. Similarly, estimating the EC-induced distortions in the DW images would be interfered by head motions. Also, the intensity inhomogeneity of the dMRI data could bias the registration process because the registration per se is driven by the intensity difference. In addition, registering the pseudo *b*_0_ image (which is distortion-free, without the intensity inhomogeneity, and in the space of the anatomical image) to the real *b*_0_ image (which is in the dMRI space) would be accurate only if the distortions and the intensity inhomogeneity of the latter are substantially mitigated. Furthermore, the diffusion tensors or the MAP-MRI coefficients for synthesizing the pseudo dMRI data should be estimated from the real dMRI data where the artefacts are reduced. Consequently, if these issues are not addressed in an integrated fashion, those not taken into consideration would degrade the performance.

Second, we develop a 1D registration algorithm which is dedicated for the susceptibility-induced distortions. Some of the previous studies (Huang et al., 2008; Irfanoglu et al., 2015) use general-purpose three dimensional (3D) registration algorithms, such as LDDMM (Beg et al., 2005; Miller et al., 2006) and Advanced Normalization Tools (ANTs) (Avants et al., 2008), to conduct the registration. Employing these 3D registration algorithms to solve the 1D distortion correction problem has at least two drawbacks. First, the calculated free-form deformations have to be further constrained to the PE direction (Irfanoglu et al., 2015), leading to unnecessary calculations. Second, which is more serious, is that the convergence of the registration may not always correspond to the convergence of distortion correction because the registration algorithms have higher degrees of freedom than the distortion problem per se. To avoid these drawbacks, we developed 1D-LDDMM with a Gauss-Newton optimization scheme for the registration algorithm, which is proved efficient and effective.

Third, we take the intensity inhomogeneity into consideration by including the bias field in the registration process. To the best of our knowledge, this is the first attempt in the field of anatomical image-based correction methods. The direct benefit of doing so is the mitigation of intensity inhomogeneity in the dMRI data. Another benefit is that the bias field could improve the stability of registration, which is crucial for the ill-posed registration problems.

### 4.2 Limitations

The proposed DACO method has some limitations. First, the method does not address the pile-up problem, which is usually encountered in regions with significant contractive distortions. In these regions, the signals from distant tissues are piled together in single voxels, therefore, in theory, these signals can be fully recovered only when the information of how the signals are piled up is known. One approach to recover the piled signals is to use two dMRI datasets which have exact the same acquisition parameters except opposite PE directions. In this approach, the regions with contractive distortions in one dMRI data would have dilative distortions in another dMRI data, therefore provides information to restore the piled signals (Andersson et al., 2003; Hedouin et al., 2017; Irfanoglu et al., 2015). Another approach is to explicitly measure the point spread function (PSF) (In & Speck, 2012; Oh et al., 2012), which represents how a point in the distortion-free space spreads in the distorted space. The distorted image can be formulated as the convolution of the undistorted image with the PSF, therefore, the piled signals could be recovered by deconvolving the distorted images with the PSF. Both approaches could properly restore the piled signals, however, the time spent on acquiring the additional images is significant. In the reverse gradients approach, the acquisition time is doubled since two duplicate sets of dMRI data are acquired, like the HCP dMRI data. In the PSF approach, the PSF is measured through an additional encoding (i.e., the PSF-encoding), which is also time consuming. Because DACO does not address the pile-up problem, for regions where the problem is significant, the results should be interpreted with caution.

Another limitation of the proposed method is that the translation of head motions confounds with the EC-induced distortions, which causes higher variations of the translation along the PE direction than other directions. Therefore, the estimated head motion parameters of the translation along the PE direction should be interpreted carefully.

Furthermore, this method corrects head motions among the DW images, however, it does not consider the situation of image signal dropout (Andersson et al., 2016). This artefact is usually encountered when the subjects have large head motions and/or when the DW images are acquired with high b-values. The corrected dMRI data should be inspected carefully in these critical situations.

The algorithm assumes that the distortion maps arise from the susceptibility are diffeomorphic. Although this assumption is adequate in the testing cases presented in the study, we cannot assure it is still valid in images suffering from very severe distortions. In these critical situations, the results should be inspected carefully because our method may fail to restore the distortions properly.

### 4.3 Future Extensions

Some issues not addressed in this study will be considered in the future. First, this study uses a static approach to estimate the susceptibility-induced off-resonance fields when the head moves. It is, however, well-known that the susceptibility-induced distortions depend not only on the static files but also on the head orientations, though the former factor accounts for a greater proportion. Anderson et al. (Andersson et al., 2018) proposed a method to estimate the orientation-dependent deviations from the static susceptibility-induced off-resonance fields using a first-order approximation. Another possible solution is suggested by Boegle et al. (Boegle et al., 2010), which estimates these deviations under a theoretical framework along with a susceptibility model of the human head, where the head model is generated from the segmentation results of MRI anatomical image such as T1w or T2w images. In the future extension, we will investigate this issue using these potential methods.

In addition, this study mitigates the head motion problem by volume-wise registration of the DW images. This approach can correct the inter-volume head movements but does not take intra-volume motions into account. Recently, some algorithms have been proposed to address intra-volume motions by conducting the registration slice-by-slice (Andersson et al., 2017). Another issue related to intra-volume motions is the signal outlier detection and signal recovery (Andersson et al., 2016). In the future extension, we will address these issues.

### 4.4 Conclusion

This study presents an integrated registration-based framework to mitigate various artefacts of dMRI data with good correction quality. The method uses the anatomical image (T1w or T2w), which is routinely acquired, to serve as the reference image and drive the correction. Therefore, the efforts to conduct this algorithm is minimum. We believe our method is beneficial to most dMRI data, particularly to those without acquiring the field map or blip-up and blip-down images.

## Acknowledgement

Data were provided [in part] by the Human Connectome Project, WU-Minn Consortium (Principal Investigators: David Van Essen and Kamil Ugurbil; 1U54MH091657) funded by the 16 NIH Institutes and Centers that support the NIH Blueprint for Neuroscience Research; and by the McDonnell Center for Systems Neuroscience at Washington University.

Data were provided [in part] by OASIS-3: Principal Investigators: T. Benzinger, D. Marcus, J. Morris; NIH P50 AG00561, P30 NS09857781, P01 AG026276, P01 AG003991, R01 AG043434, UL1 TR000448, R01 EB009352. AV-45 doses were provided by Avid Radiopharmaceuticals, a wholly owned subsidiary of Eli Lilly.

## Appendix A

Let ***φ***_*ϵ*_ be a family of diffeomorphisms parameterized by the real variable *ϵ*, such that ***φ***_0_ = ***φ*** and

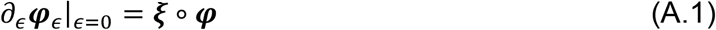

From this definition, we can show that the variation of D***φ**_ϵ_* is

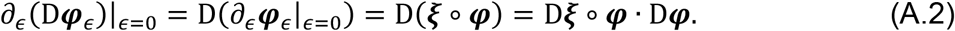

Therefore, the variation of |D***φ**_ϵ_*| is

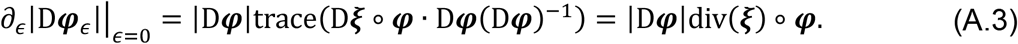

Since 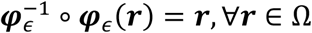, we have

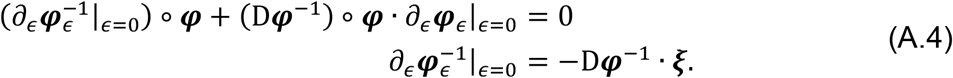

Furthermore, 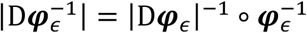, its variation is

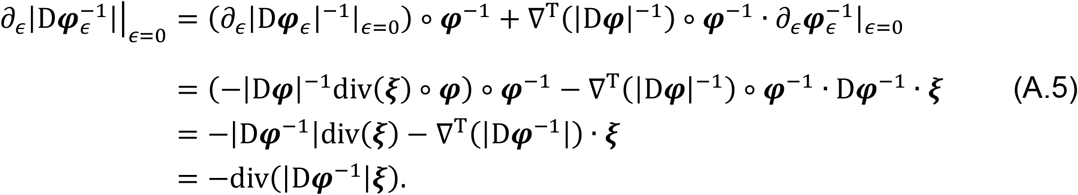

The action of ***φ*** on an image *f*_0_ is defined as ***φ*** * *f*_0_ = |D***φ***^−1^|*f*_0_ ∘ ***φ***^−1^, so

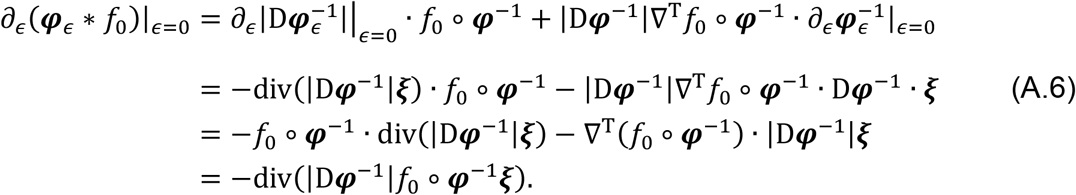

The data matching functional between images *f*_0_ and *f*_1_ is defined as

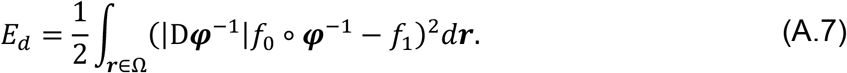

The first variation of *E_d_* in the direction *ξ* is computed via

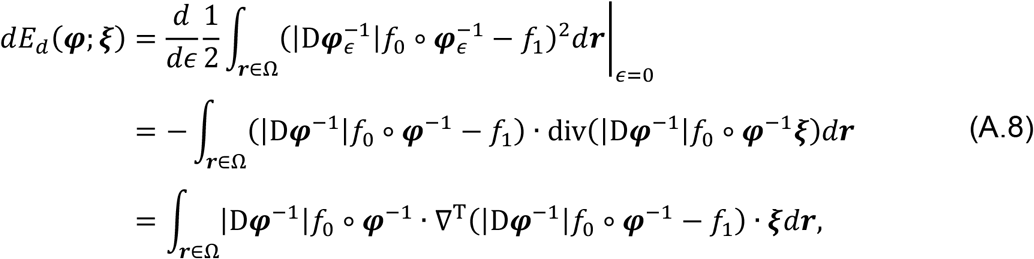

where the last line uses the divergence theorem, assuming that *ξ* vanishes on *∂*Ω, or is tangent to *∂*Ω, or vanishes at infinity. Therefore, the first Gâteaux differential of *E_d_* with respect to ***φ*** is

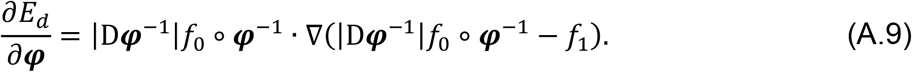

This rephrases equation (9.29) in (Younes, 2019). The second variation of *E_d_* in directions *ξ*_1_ and *ξ*_2_, with the positive definite approximation, is

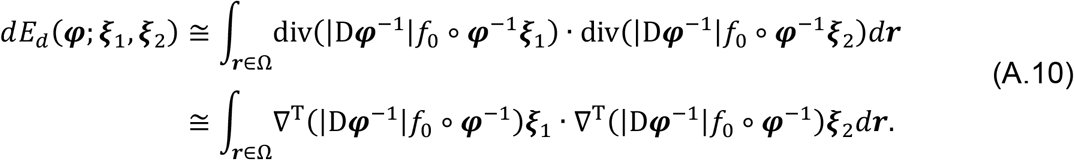

Therefore, the approximated second Gâteaux differential of *E_d_* with respect to ***φ*** is

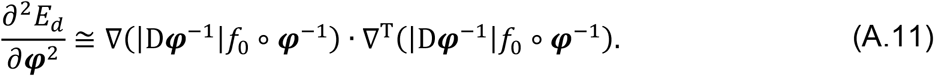

Under the LDDMM framework, the first derivative of *E_d_* with respect to the initial velocity ***u*** is expressed as (Younes, 2019)

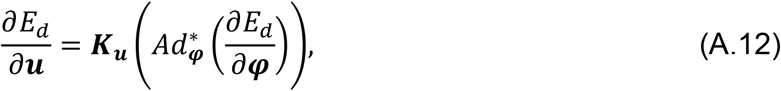

where 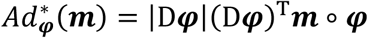. With explicit elaboration, we obtain

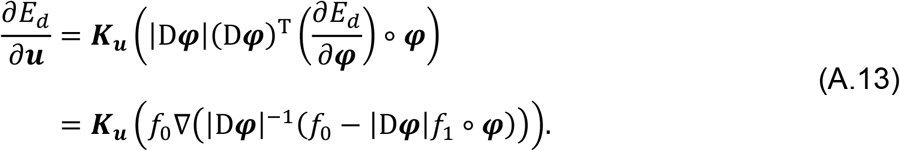

Unlike the first derivative, there is no explicit formulation between the second derivatives. This is not a serious problem to us, since we only need a reasonable approximation which suffices to conduct the Gauss-Newton optimization scheme. Observing Eq. (A13), *Ad*\* transports 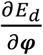, which situates at ***φ***^−1^, to the identity. We employ similar operation to approximate the second derivative of *E_d_* with respect to ***u***, yielding

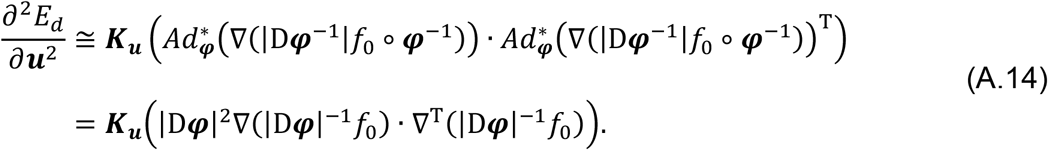

